# CleanFinder: A Scalable Framework for Comprehensive Genome Editing Analysis

**DOI:** 10.1101/2025.10.23.684080

**Authors:** H. Ramachandran, J. Dobner, T. Nguyen, S. Binder, I. Tolle, I. Vykhlyantseva, J. Krutmann, A. Miccio, C. Staerk, M. Brusson, Z. Kontarakis, A. Prigione, A. Rossi

## Abstract

Genome editing experiments routinely generate complex mixtures of alleles rather than single, predefined outcomes. Resolving these heterogeneous edits across diverse editing modalities, sequencing platforms, and multiplexed designs remains a persistent analytical challenge. To address this, we developed CleanFinder, a browser-native framework for genotyping genome editing outcomes using a constrained semi-global alignment strategy. Context-aware alignment modes support a broad spectrum of editing scenarios, including indels, base substitutions, and complex prime editing modifications, across nuclear and mitochondrial targets. Additional modules include an optional turbo mode for high-throughput heuristic alignment in exploratory workflows, and an allele-aware module that leverages heterozygous SNPs to detect allelic dropout. To evaluate scalability and practical performance, we applied CleanFinder to a primary small-molecule screen of 1,849 compounds in HEK293T cells. The software efficiently processed the dataset, enabling high-throughput comparison of editing outcomes and nomination of candidate compounds for follow-up analysis. Together, CleanFinder provides a flexible and scalable platform for genome editing analysis, enabling detailed genotyping and systematic comparison of editing outcomes across diverse edit types and genomic contexts.

## Introduction

The CRISPR/Cas system has revolutionized biological research by enabling precise genome manipulation with unprecedented ease and efficiency (Le Cong et al., 2013, Mali et al., 2013). Advanced programmable genome editing technologies, including CRISPR-based Base Editing (BE) and the more recently developed Prime Editing (PE), have substantially expanded the scope of precise nuclear genome engineering (Gaudelli et al., 2017; Anzalone et al., 2019). In parallel, CRISPR-free systems such as RNA-free DddA-derived cytosine base editors (DdCBE) have enabled targeted mitochondrial DNA (mtDNA) editing, further broadening the landscape of programmable genetic manipulation (Mok et al., 2020). Despite these advances, the detection and classification of editing events, including insertions and deletions (indels), complex rearrangements, and off-target effects, often require specialized computational expertise and access to advanced bioinformatics tools. This creates a bottleneck for many laboratories, particularly those who are not using genome editing approaches on a regular basis.

Existing tools for analyzing CRISPR/Cas editing data typically require local installation and command-line execution, often distributed via Python environments, containerized solutions such as Docker, or integrated into workflow managers like Nextflow (Clement et al., 2019; Amit et al., 2021; Sanvicente-García et al., 2023; Di Tommaso et al., 2017). While powerful, these approaches present practical barriers for users without computational expertise and can complicate rapid exploratory analyses. Web-based implementations of some of these tools (Clement et al., 2019, Park et al., 2017) provide greater accessibility but are often constrained in throughput, for example by limiting the number of samples processed per run or supporting only single-sample analysis.

In addition, web-based tools are often designed around short-read Illumina genotyping workflows (Schmid-Burgk et al., 2014), with only partial adaptation to the distinct error profiles and read-length characteristics of long-read platforms such as Oxford Nanopore Technologies (ONT) (Park et al., 2017, Nguyen et al., 2022).

To address these analytical and practical limitations, we developed CleanFinder as a modular framework for high-resolution genotyping of genome editing outcomes. Built upon a constrained semi-global alignment strategy, the platform supports context-aware classification across diverse editing modalities, including Cas9, Cas12, BE, and PE, and is compatible with both Illumina and ONT sequencing technologies. CleanFinder integrates dedicated analytical modules for inference of DNA repair pathway signatures through Rational Indel Meta-Analysis (RIMA) and Repairome analysis (Taheri-Ghahfarokhi et al., 2018; Lopez de Alba et al., 2025), together with allele-aware classification based on heterozygous SNPs to detect allelic dropout. In addition, the dedicated PE mode performs competitive wild-type versus intended-edit scoring to resolve complex insertion, deletion, and substitution events across multiple loci and cellular contexts. A full comparison of these features against existing platforms is summarized in **Table S1**.

To accommodate both exploratory and large-scale workflows, CleanFinder is available both as a browser-native web interface and as a full-feature command-line implementation. In addition to the standard deterministic pipeline, the platform implements a high-throughput heuristic Turbo mode for rapid analysis of standard editing experiments. The command-line interface (CLI) version reproduces the full analytical functionality of the web application, including context-aware alignment, editing classification, RIMA/Repairome profiling, and allele-aware dropout detection, while eliminating browser memory limitations. This enables scalable batch processing of large datasets and seamless integration into custom bioinformatics pipelines. An integrated genome viewer provides locus-level visualization of gene architecture, coding regions, translated protein sequence, and optional gRNA placement, facilitating contextual interpretation of editing outcomes within their genomic and functional framework.

We demonstrate the scalability and practical applicability of CleanFinder in a high-throughput small-molecule screen comprising 1,849 compounds in HEK293T cells, enabling systematic comparison of prime editing outcomes across experimental conditions.

Together, CleanFinder establishes a flexible and scalable analytical framework for systematic comparison of genome editing outcomes across platforms, loci, and experimental designs.

## Results

### CleanFinder: a modular framework for genome editing outcome analysis

The modular architecture of CleanFinder is summarized in **Figure 1**. The platform is organized into four functional layers: user-defined options, processing logic, analytical outputs, and auxiliary modules. Input parameters include organism or reference locus, optional gRNA information, and customizable alignment settings. Sequencing reads derived from any major NGS platform (Illumina, PacBio, or ONT) are automatically parsed and directed to biologically informed analysis modes (e.g., Cas9/Cas12, BE, or PE). A default high-resolution alignment workflow is complemented by a heuristic turbo mode for exploratory and fast analyses.

**Figure 1.**
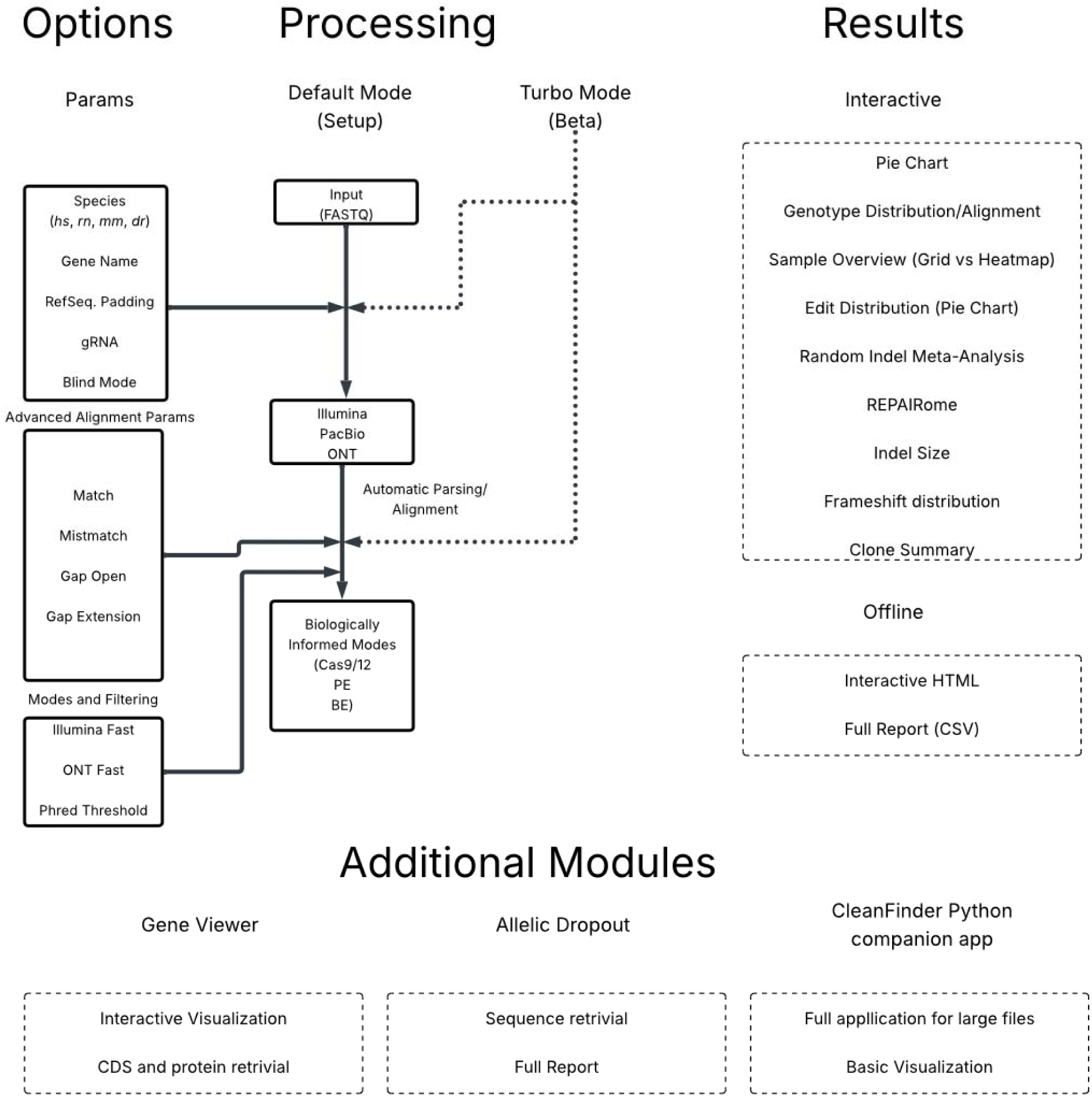
Modular architecture of CleanFinder. The CleanFinder platform is organized into modules that ar structured into four functional layers: Options, Processing, Results, and Additional Modules. The Options layer allow specification of target organism (species), reference locus/gene name, gRNA sequence, alignment parameters (match, mismatch, gap penalties), and sequencing-specific filters. In the Processing layer, FASTQ inputs (Illumina, PacBio, or ONT) undergo automated parsing and amplicon detection, followed by biologically informed mode selection (e.g., Cas9/Cas12, Prime Editing, Base Editing). A default high-resolution alignment workflow is complemented by a heuristic high-throughput mode for rapid exploratory analyses. The Results layer provides interactive summaries of genotype distributions, edit-type composition, indel spectra, frameshift status, and repair pathway inference through Rational Indel Meta-Analysis (RIMA) and Repairome signatures. Output is available as interactive HTML reports or structured CSV files. Additional modules include an integrated genome viewer for gene-level contextualization, an allele-aware dropout module based on heterozygous SNP detection, and a Python companion application enabling batch processing of large datasets.

The results layer provides structured summaries of editing outcomes (**Figure 2**), including genotype distributions, indel spectra, frameshift classification, and repair pathway inference via RIMA and Repairome signatures (*Wimberger et al 2023, Lopez de Alba et al., 2025*). Outputs are accessible through interactive visualization and exportable reports. Additional modules extend interpretability through allele-aware dropout detection and an integrated genome viewer that contextualizes editing events within gene structure. A companion Python implementation enables scalable batch processing.

**Figure 2.**
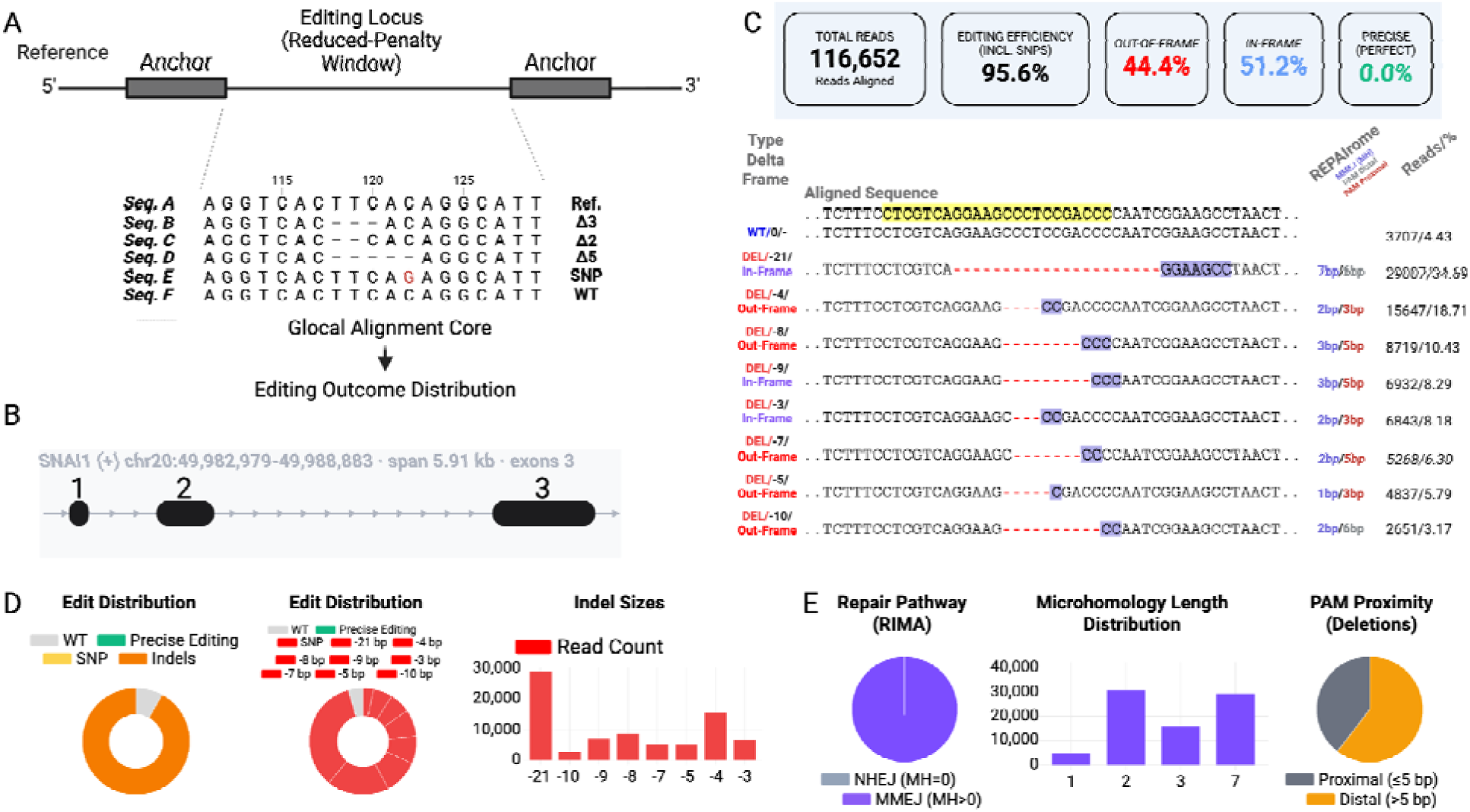
**Anchor-guided alignment and structured classification of Cas12 editing outcomes in human induced pluripotent stem cells**. Analysis of a Cas12 RNP-mediated genome editing experiment targeting SNAI1 in human iPSCs is exemplarily shown to illustrate the CleanFinder workflow. (A) Constrained semi-global (“glocal”) alignment strategy. High-confidence 5′ and 3′ anchor sequences define a reduced-penalty window centered on the editing locus. Within this window, semi-global alignment preserves full-length read continuity while accommodating insertions, deletions, and substitutions. (B) Integrated Genome Viewer module displaying the targeted *SNAI1* locus with exon organization and transcript structure, enabling contextual interpretation of editing events. (C) Structured outcome classification and summary metrics, including total aligned reads, overall editing efficiency, frame statu (in-frame versus out-of-frame), and precise editing frequency. Representative aligned sequence illustrate deletion size and frame annotation together with Repairome labeling. (D) Quantitative distributions of editing outcomes, including global edit composition and size-resolved indel spectra. (E) Repair pathway inference through Rational Indel Meta-Analysis (RIMA), microhomology length distribution, and PAM proximity analysis, enabling characterization of repair dynamics. (B) - (E) are representative of the visualizations provided by CleanFinder. They have been recreated for high-resolution rendering within this article.

In the user-facing analysis setup, the users define the reference sequence or genomic locus, specify analysis windows (e.g., gRNA-centered anchor regions), upload sequencing reads, and configure alignment and filtering parameters prior to execution. This design supports both exploratory experimentation and standardized high-throughput workflows (cleanfinder.org).

### Anchor-Guided Semi-Global Alignment for High-Resolution Genotyping

CleanFinder employs a constrained semi-global alignment strategy (**Figure 2A**), a dynamic programming approach in which terminal gaps are not penalized (Durbin et al., 1998). Classical local alignment approaches such as Smith–Waterman are optimized to identify the highest-scoring subsequence between two sequences, but may truncate flanking context and fragment complex editing events (Smith et al., 1981). Conversely, strict global alignment enforces end-to-end matching and does not naturally accommodate terminal overhangs, which are common in long-read sequencing data (Needleman et al., 1970).

To enable accurate classification of diverse editing outcomes across sequencing platforms, CleanFinder employs a constrained semi-global (“glocal”) alignment strategy (Durbin et al. 1998). Two high-confidence anchor sequences define a reduced-penalty window centered on the editing locus. Within this window, reads are aligned using semi-global alignment without penalizing terminal gaps. This anchor-guided design restricts alignment to the biologically relevant region of interest while preserving full-length read continuity, enabling accurate reconstruction of insertions, deletions, substitutions, and composite editing events across sequencing platforms.

To illustrate the analytical capabilities of CleanFinder, we analyzed a Cas12 RNP-mediated editing experiment targeting *SNAI1* in human induced pluripotent stem cells (iPSCs).

Figure 2B illustrates the integrated Genome Viewer module. Editing outcomes can be visualized in the context of gene architecture, including exon organization, strand orientation, and coding regions. This enables direct interpretation of editing events relative to transcript structure and facilitates assessment of potential coding impact.

Aligned reads are then classified according to edit type, size, and coding consequence. As illustrated in Figure 2C, CleanFinder reports total aligned reads, overall editing efficiency (including SNPs), precise editing frequency, and effect on the coding triplets (in-frame versus out-of-frame). Representative aligned sequences demonstrate systematic annotation of deletion size and reading-frame impact, together with Repairome labeling of repair characteristics.

Quantitative summaries of editing distributions are provided in Figure 2D, including global edit composition and size-resolved indel spectra. Finally, Figure 2E presents repair pathway inference through Rational Indel Meta-Analysis (RIMA), microhomology length distribution, and PAM proximity analysis, enabling mechanistic interpretation of DNA repair patterns.

### Cross-Platform Validation Across Diverse Nuclease Systems

To evaluate CleanFinder across distinct CRISPR nucleases and editing strategies, we analyzed Cas9- and Cas12-mediated genome editing experiments in human iPSCs, encompassing both knockout (KO) and knock-in (KI) designs (**Figure S1, S2**). Across both Cas9 and Cas12 systems, CleanFinder consistently resolved deletion size, frame status, microhomology length, and PAM proximity. Heatmap visualizations demonstrate genotype quantification across independent clones and editing efficiency (**Figure S1A**). Representative aligned sequences illustrate MMEJ-associated deletions in *VHL* and *DMD* knockout experiments, with locus-dependent differences in microhomology usage and PAM-proximal versus PAM-distal deletion patterns that reflect nuclease cleavage architecture. In contrast, knock-in experiments at the *LRPPRC* and *XPA* loci showed precise homology-directed repair (HDR) signatures with accurate reconstruction of intended integrations and minimal indel-associated microhomology (**Figure S1B**). These analyses confirm that CleanFinder supports structured classification of genome editing outcomes across nuclease systems and distinguishes NHEJ/MMEJ-driven knockout events from HDR-mediated knock-in designs in iPSCs.

### Benchmarking Consistency on Illumina and Oxford Nanopore Technologies

Next, we benchmarked CleanFinder performance across Illumina and ONT sequencing data, focusing on genotype concordance in both single-clone and bulk-edited samples. ONT reads were basecalled using the super accuracy (SUP) model, as lower stringency modes (fast and high accuracy) may introduce elevated indel noise impairing reliable genotype resolution. As shown in **Figure S2A**, genotype frequencies derived from ONT exhibited strong concordance with Illumina measurements in single-clone analyses across multiple genes and dozens of independently derived clones (Pearson r = 0.97, p < 0.001). Similarly, bulk-edited samples demonstrated high cross-platform agreement (**Figure S2B**; Pearson r = 0.9, p < 0.001). Despite this overall concordance, isolated discrepancies were observed especially in homopolymeric regions of ONT reads in line with the literature (Wang Liu-Wei et al., 2023). An example is highlighted for a genetically modified *HIF1A* clone in HEK293T cells (**Figure S2A**), where indel miscalling occurred within a poly-A stretch. Similar homopolymer-associated indel slippage events were detected at additional loci containing repetitive nucleotide stretches (data not shown).

Importantly, these localized discrepancies did not substantially impact global genotype frequency estimates. When basecalled in SUP mode, ONT data showed high concordance with Illumina measurements, supporting the use of CleanFinder for long-read genotype analysis despite known ONT-specific indel biases (Wick et al., 2019). Interpretation of individual genotypes within homopolymeric regions should therefore be performed with caution.

To evaluate analytical sensitivity across sequencing platforms, we performed controlled dilution experiments at the *HOXB7* locus by mixing validated knockout (Δ13) amplicons with wild-type (WT) DNA at defined ratios spanning 0–100%. This experimental setup enabled precise assessment of CleanFinder’s ability to detect and quantify indel alleles across a wide range of frequencies using long-read sequencing (Oxford Nanopore Technologies; ONT) (**Figure S3A**). In parallel, analytical sensitivity under short-read conditions was assessed using *in silico*–generated Illumina-based datasets composed of defined proportions of WT and Δ10 sequences. Low-frequency KO events (0.02%, 0.1%, 1%, and 10%) were simulated and compared to the observed allele frequencies prior analysis (**Figure S3B**).

Together, these approaches demonstrate robust detection across a broad dynamic range (0.02–100%) using both experimentally generated (ONT) and computationally simulated (Illumina) data.

### k-mer–guided pre-alignment filtering improves computational efficiency

To reduce computational overhead during read alignment, we implemented a k-mer–guided pre-alignment filter that restricts full semi-global alignment to reads containing short exact matches to the target locus (**Figure S4**). As illustrated schematically (**Fig. S4A**) , reads lacking the required k-mer seed are discarded prior to dynamic programming alignment, whereas candidate reads proceed to full classification. This reduces computational load by limiting alignment to locus-relevant reads.

Benchmarking demonstrated a substantial increase in processing speed for both Illumina and ONT datasets (**Fig. S4B**). For Illumina reads, median throughput increased from approximately 8.4 thousand reads per second (normal mode) to 18 thousand reads per second (fast mode).

For ONT datasets, throughput increased from approximately 0.5 thousand reads per second to 1.89 thousand reads per second. The magnitude of speed improvement was therefore particularly pronounced for long-read data.

These values represent empirical benchmarks under representative conditions. Processing speed depends on dataset characteristics, including read length, file size, and, critically, the proportion of reads unrelated to the targeted locus. In multiplexed or off-target–rich datasets, the k-mer filter discards unrelated reads early in the workflow, leading to disproportionately greater acceleration.

### Robust Classification of Base Editing and Complex Prime Editing Events

To extend validation beyond double-strand break–based genome editing, we next evaluated CleanFinder in base editing experiments, which introduce precise nucleotide substitutions without generating double-strand breaks (Gaudanelli et al., 2017, Mok et al., 2020, Antoniou et al., 2021). Human iPSC models enable high-throughput compound screening; however, screening for modifiers of editing efficiency requires accurate and scalable heteroplasmy quantification (Zink et al., 2026). To evaluate CleanFinder in this context, we used mtDNA base editing in human induced pluripotent stem cells (iPSCs) to introduce a pathogenic *MT-ND5* mutation associated with Leigh syndrome and subsequently analyzed the editing outcomes (Figure 3A). Unlike nuclear CRISPR editing, mitochondrial base editing generates heteroplasmic nucleotide conversions within multi-copy mitochondrial genomes (Mok et al., 2020; Rossi et al., 2022). We analyzed five independent edited iPSC clones using ONT long-read sequencing (∼2 kb amplicons). CleanFinder accurately quantified G→A conversion and heteroplasmy levels, leveraging anchor-guided alignment and k-mer–assisted read filtering to efficiently process full-length reads. Estimated heteroplasmy frequencies closely matched values obtained using the dedicated mitochondrial analysis pipeline Mitopore (Dobner et al., 2024), demonstrating concordant quantification of mtDNA base editing outcomes (Figure 3B). We next evaluated nuclear CRISPR-based adenine base editing (ABE) in human hematopoietic stem and progenitor cells (HSPCs) targeting the *HBG1/2* γ*-globin* promoters locus, a clinically relevant site for fetal hemoglobin modulation, to induce programmable T→C nucleotide conversion (Figure 3C). CleanFinder accurately detected targeted nucleotide conversions within defined editing windows and distinguished intended base substitutions from bystander edits and residual indel events. Substitution frequencies were robustly quantified across samples, demonstrating applicability in clinically relevant primary cell types. We additionally performed an in silico simulation of cytosine base editing (CBE) across defined editing windows. CleanFinder reliably detected expected C→T conversions and distinguished single from clustered substitutions, confirming robust substitution-based genotype resolution (Figure 3D).

**Figure 3.**
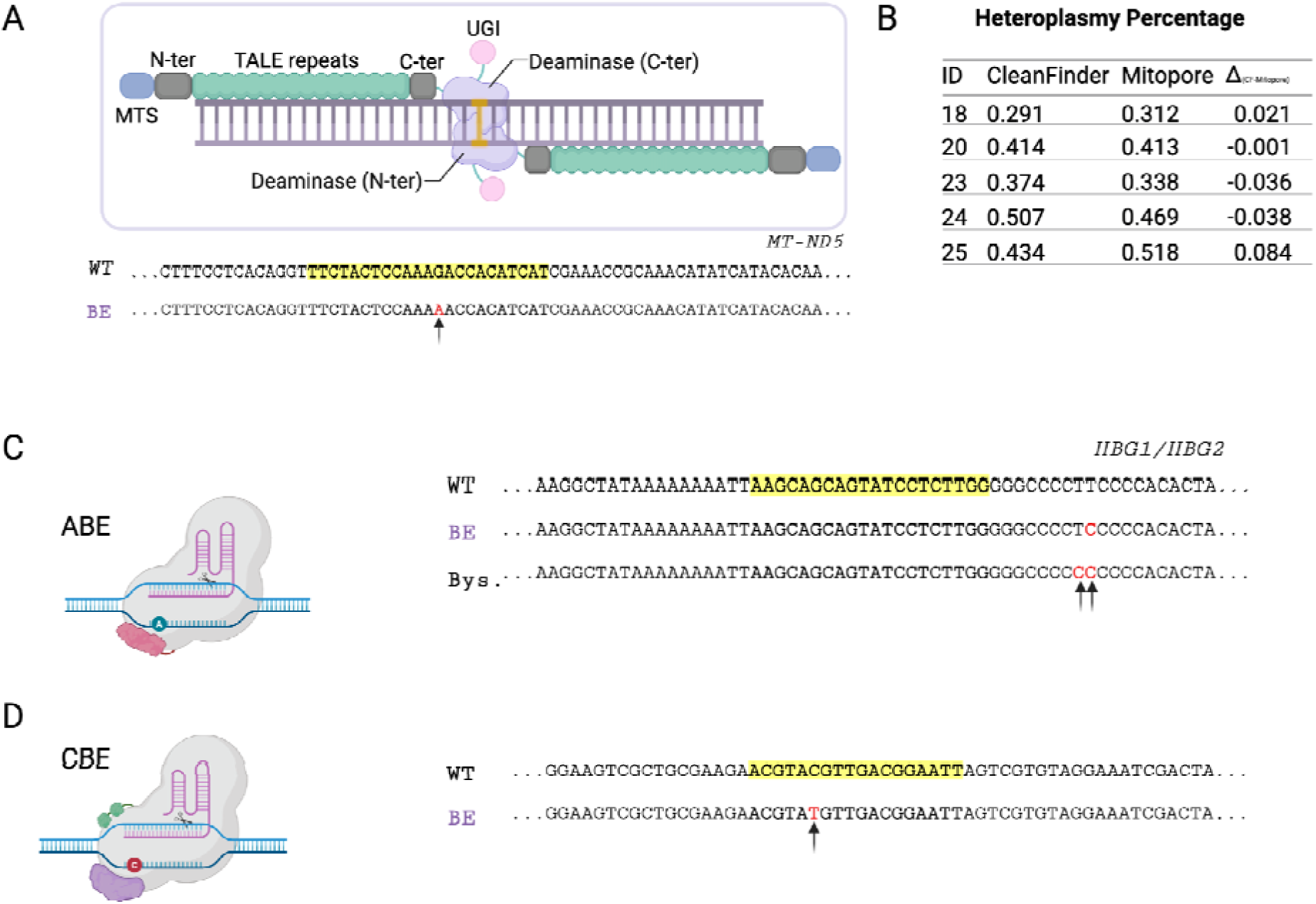
Validation of CleanFinder in base editing applications. (A, top): Schematic representation of TALE-based mitochondrial base editing in human iPSCs. Deaminase domains induce targeted nucleotide conversion within mtDNA without double-strand breaks. (A, bottom): Analysis of mtDNA base editing outcomes (G→A conversions) from Oxford Nanopore long-read (∼2 kb) amplicons. (B) Quantification of heteroplasmy levels across five independently edited iPSC clones (IDs 18, 20, 23, 24, 25). CleanFinder estimates heteroplasmy percentages using anchor-guided semi-global alignment and k-mer–assisted read filtering. Editing frequencies show close concordance with values obtained using the dedicated mitochondrial analysis pipeline Mitopore. (C): Nuclear CRISPR-based adenosine base editing (ABE) T→C conversions in human hematopoietic stem and progenitor cells (HSPCs), illustrating precise nucleotide substitution within defined editing windows. (D): In silico simulation of cytosine base editing (CBE) demonstrating detection of C→T conversions and clustered substitution events within programmable editing windows.

Prime editing (PE) enables programmable introduction of precise nucleotide substitutions, insertions, and deletions without relying on co-transfection of donor templates (Anzalone et al., 2019, Chen et al., 2023, Binder et al., 2024). Depending on configuration, PE can operate in a nickase-based format (e.g., PE2/PE3) or in nuclease-enabled forms that increase editing efficiency but also generate more complex allelic architectures (Adikusuma et al., 2021, Peterka et al., 2022). These configurations frequently produce heterogeneous outcomes, including partial edits, insertion–substitution combinations, short indels, or shifted integration boundaries that complicate alignment and classification.

To handle more complex PE outcomes, such as difficult-to-align insertions arising from local mismatches, short indels, or integration site shifts, CleanFinder implements a dual competitive reference scoring framework (Figure 4A). Each read is aligned against both the wild-type (WT) reference and the intended prime-edited (PE) reference. Classification is determined by comparative alignment scoring, enabling robust discrimination between WT alleles, precisely edited PE alleles, and intermediate or bystander variants.

**Figure 4.**
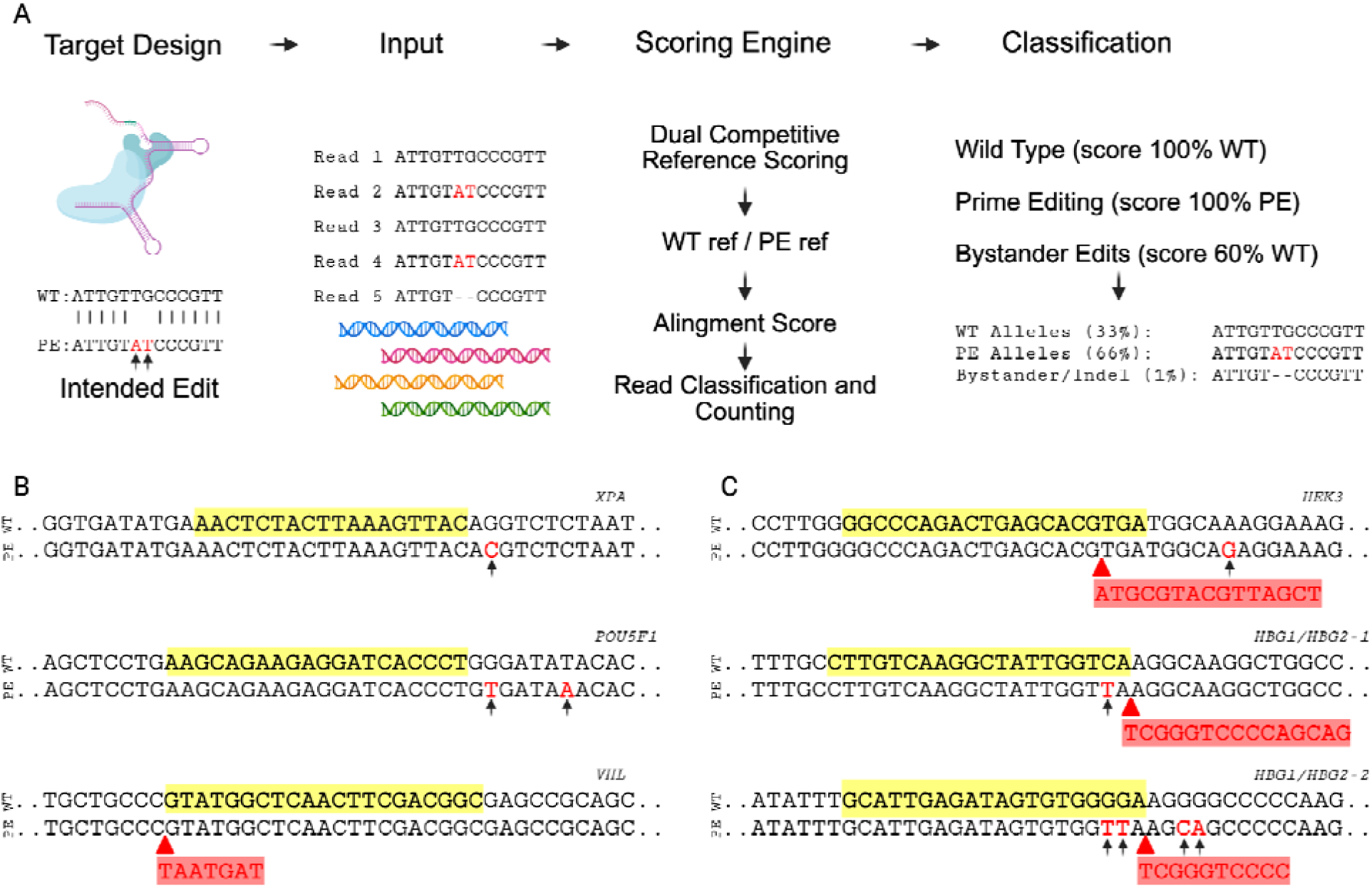
Resolution of complex prime editing outcomes across iPSCs and hematopoietic stem cells. (A, top) Schematic overview of dual competitive reference scoring for prime editing analysis. Sequencing reads are aligned against both wild-type (WT) and intended prime-edited (PE) reference sequences. Comparative alignment scoring enables classification into WT, precise PE, or intermediate alleles. (B) Prime editing experiments in human induced pluripotent stem cells (iPSCs) targeting *XPA*, *POU5F1*, and *VHL*. Substitution-only and short insertion designs are shown. Intended nucleotide change and inserted sequences are highlighted. (C) Prime editing at the *HEK3* locus in K562 cells and at the *HBG1/2* γ*-globin (*target 1 and 2) locus in in K562 cells. Increasing edit complexity is illustrated, including longer programmable insertions (*HEK3*) and composite substitution–insertion–deletion architecture (*HBG1/2* γ*-globin,* target 1 and 2). Highlighted regions indicate intended edits; red annotations denote inserted sequences or nucleotide substitutions relative to WT.

We first examined substitution-only designs at the *XPA* and *POU5F1* loci in human iPSCs (Figure 4B). These edits introduced one or two defined nucleotide substitutions within the programmable window, which were resolved at single-nucleotide resolution and clearly distinguished from wild-type alleles. We then analyzed insertion-based prime editing at the *VHL* locus in iPSCs (Figure 4B), where a short, defined sequence was integrated. The inserted sequence, highlighted within the aligned reads, was reconstructed in full within edited alleles.

Edit complexity increased further at the HEK3 locus in K562 cells (Figure 4C), where a longer programmable insertion was introduced. Full-length insertions were separated from incomplete or mis-integrated products through comparative alignment against wild-type and edited reference sequences. Finally, composite genome engineering at the *HBG1/2* γ*-globin* promoters in in K562 cells generated a spectrum of substitutions, insertions, and localized deletions within the same programmable regulatory region (Figure 4C). These multi-component architectures were resolved as discrete edited alleles distinct from wild-type reads. Collectively, these examples illustrate accurate detection of substitution-only, insertion-only, and multi-component prime editing outcomes across increasing architectural complexity and diverse cellular contexts. We found the dual-reference scoring framework to be critical for reliably distinguishing wild-type, precisely edited, and composite alleles.

### Primary Small Molecule Screen Demonstrates Scalable Analysis of Prime Editing

To demonstrate applicability of the CleanFinder framework for compound screening, a total of 1,849 compounds from the Selleck DNA Damage/DNA Repair library (L4600) and the Sigma LOPAC1280 collection were screened at 10 µM in a single-dose primary format in HEK293T cells, in line with established screening frameworks (Rothenaigner et al., 2021, Mariusz Butkiewicz et al., 2017) and prior implementations in this cell type (Li et al., 2019; Du et al., 2020; Maji et al., 2020).

Two pegRNA designs were used: a single nucleotide substitution at the *EMX1* locus and a trinucleotide insertion at the *HEK3* locus (Figure 5A). Prior to screening, timing experiments established that editing became detectable approximately 12 hours post-transfection, with extended exposure resulting in higher efficiencies (**Figure S5A, B**). Compounds were therefore added at the time of transfection to coincide with early editing events.

**Figure 5:**
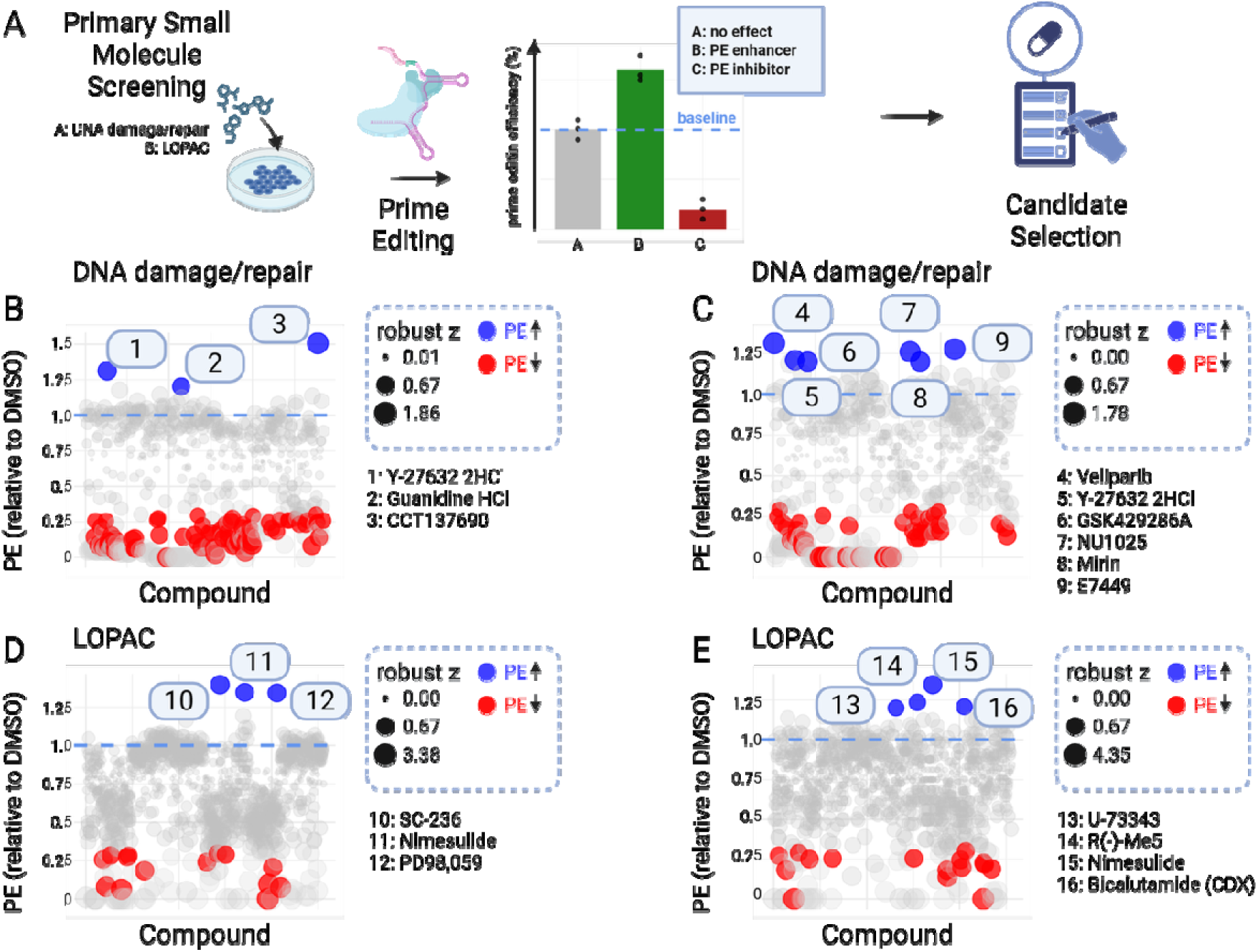
Primary small molecule screening analyzed using CleanFinder. (A) Schematic overview of the primary screening workflow. HEK293T cells were transfected with pegRNAs targeting *EMX1* (single-nucleotide substitution) or *HEK3* (trinucleotide insertion). Compounds from the Selleck DNA Damage/DNA Repair (L4600) and Sigma LOPAC1280 libraries were added at the time of transfection.

Although editing efficiencies peaked around 72 hours, several compounds showed signs of cytotoxicity or reduced transfection quality by 48 hours, as assessed by fluorescence microscopy (**Figure S5C**). To minimize confounding effects, compound exposure was thus limited to 48 hours. Cell morphology was monitored visually, and cytotoxicity was further evaluated by nuclear staining following Cas9-based transfection and quantified using Fiji image analysis (**Figure S5D**).

Editing efficiency was quantified relative to DMSO controls and ranked using robust z-score analysis. (B–C) DNA damage/repair library. Scatter plots showing Prime Editing efficiency relative to DMSO for individual compounds. Blue points indicate compounds associated with increased PE efficiency relative to baseline; red points indicate reduced efficiency. Circle size reflects robust z-score magnitude. Numbered compounds correspond to prioritized molecules listed beside each panel. (D–E) LOPAC library analyzed under identical conditions. Distribution patterns illustrate that most compounds cluster near baseline, with a subset showing measurable deviations. Dashed lines indicate DMSO baseline levels. Results represent observations from a primary screen and were not subjected to secondary validation within this study.

Editing efficiencies were quantified using CleanFinder, normalized to DMSO-treated controls, and ranked using robust z-score analysis. Across both libraries, most compounds clustered near baseline levels (Figure 5B**–E****, Figure S5E**). A subset exhibited positive or negative deviations in PE efficiency relative to control-treated samples. Among compounds showing upward deviations were Y-27632 2HCl, Guanidine HCl, CCT137690, Veliparib, GSK429286A, NU1025, Mirin, and E7449 (DNA damage/repair library), and SC-236, Nimesulide, PD98,059, U-73343, R(–)-Me5, and Bicalutamide (CDX) (LOPAC library). The magnitude of observed effects was generally moderate and varied moderately between substitution and insertion designs (**Figure S5E**). Compounds exceeding predefined robust z-score thresholds above 2 were nominated as candidate modulators for future validation.

Based on effect magnitude, reproducibility across editing designs (substitution and insertion), and prior reports suggesting context-dependent effects of histone deacetylase (HDAC) inhibition on PE efficiency (Liu et al., 2022), selected HDAC inhibitors were prioritized for secondary dose–response analysis. Secondary screening confirmed the inhibitory effect of selected HDAC inhibitors on prime editing efficiency for both substitution and insertion designs at the *EMX1* and *HEK3* loci, respectively (data not shown).

Furthermore, dose–response analysis of insertion-based prime editing at two independent loci (*HEK3* and *VHL*) demonstrated reproducible, sigmoidal modulation of editing efficiency (**Figure S6**). Editing efficiencies exhibited moderate dose-dependent decreases with increasing inhibitor concentrations, and fitted four-parameter logistic models yielded ED50 values in the low micromolar range. These findings indicate that CleanFinder reliably captures graded, concentration-dependent editing responses across distinct genomic targets.

While confirmatory validation was beyond the scope of this study, primary screening datasets provide quantitative insight into experimental variability and effect magnitude and are commonly disseminated to support downstream prioritization efforts (*Butkiewicz et al., 2017, Inglese et al., 2007*). The observations presented here intend to illustrate screening-scale performance of the analytical framework, not hit validation *per se*. Collectively, these findings demonstrate consistent quantification and ranking of editing outcomes across substitution and insertion designs in a screening-scale setting.

Beyond its application to screening-scale datasets, CleanFinder was designed as a modular framework supporting multiple deployment modes and analytical extensions. We therefore next describe additional implementation layers that extend the core browser-based application.

## CleanFinder: Additional Modules

### CleanFinder command-line integration (CLI)

To support high-throughput and reproducible processing of large sequencing datasets, we implemented CleanFinder as a Python-based command-line interface (CLI) (**Figure S7A–C**). The CLI version implements the full CleanFinder analysis workflow and exposes core parameters (e.g., mode selection, anchor length, mismatch tolerance, quality filtering, and gRNA-centered windowing) for integration into scripted pipelines and batch execution. In addition to reporting genotype distributions and frame status, the CLI outputs aligned allele representations and computes deletion-associated microhomology features to summarize repair pathway usage (e.g., MMEJ versus NHEJ) where applicable (**Figure S7C**). These outputs are designed to preserve interpretability and support downstream automation through structured exports (e.g., CSV files containing aligned reference/read strings).

### Turbo Mode

In parallel, we developed CleanFinder Turbo, a browser-native module designed specifically for rapid screening and interactive exploration (**Figure S7D–E**). CleanFinder Turbo uses a heuristic read-localization strategy: in this context, heuristic means that read placement and classification are determined using fast, rule-based approximations (anchor detection with bounded mismatches) rather than computing an optimal dynamic-programming alignment for every read. Conceptually, this mirrors the broader class of seed/anchor-based heuristic strategies used to accelerate sequence mapping in general-purpose tools such as BLAST and short-read aligners (Altschul et al., 1990). Practically, Turbo extracts forward anchors from the reference and computes reverse-complement anchors to enable strand-agnostic detection. Reads are screened by mismatch-bounded anchor scanning, and classification proceeds from the extracted internal segment and length delta relative to the expected reference middle region. This design minimizes per-read computation and enables processing rates on the order of ∼10^5 reads per second, making Turbo well-suited for primary screening and multi-well plate-scale quality control without the full alignment detail, repair pathway inference, or advanced visualization modules available in the main application.

### Allelic Dropout

On-target large deletions are a well-documented and undesirable outcome of genome editing (*Kosicki et al., 2017; Simkin et al., 2022*). In addition to quantifying editing outcomes, CleanFinder incorporates a dedicated module to detect allelic dropout (Figure 1**, Figure S8A, B**), a common artifact in CRISPR experiments in which one allele is preferentially lost due to large deletions or primer-site mutations.

The dropout detection strategy leverages naturally occurring heterozygous single nucleotide polymorphisms (SNPs) within the amplicon as endogenous genomic barcodes. In a heterozygous sample, the presence of a biallelic SNP provides evidence that both parental alleles were successfully amplified. Conversely, failure to detect a known heterozygous SNP indicates a monoallelic state consistent with allelic dropout. Critically, the use of long-read sequencing is often necessary to resolve the segregation and phasing of these markers across the entire locus. This approach was applied, for instance, to a 3 kb amplicon encompassing the *XPA* locus, where long-read sequencing was required to link a homozygous deletion with a flanking heterozygous SNP (**Figure S8C**). By phasing these distant genomic features on single molecules, CleanFinder confirmed the maintenance of diploidy and verified balanced heterozygous representation (Papadopoulou et al., 2025). The final output provides a definitive classification of “BIALLELIC” or “MONOALLELIC,” together with coverage and allele frequency metrics for the highest-confidence SNP.

The allelic dropout module was benchmarked on both synthetic datasets and biological loci with known allelic configurations, as well as on whole-genome sequencing (WGS) data aligned using minimap2 (Li et al., 2018). Across these evaluations, the long-read phasing workflow demonstrated high sensitivity for detecting allelic imbalance and large deletions while preserving haplotype context. These results indicate that the allelic dropout module, when combined with long-read phasing, enables robust detection of large deletions together with their allelic assignment. In contrast to indel-centric workflows that quantify deletion frequencies alone, this approach explicitly evaluates allelic representation and can reveal imbalance events that would remain undetected in conventional editing analyses.

## Discussion

Genome editing experiments increasingly generate heterogeneous allelic architectures that extend beyond “simple” insertion–deletion events at a defined cut site. Composite indel–substitution events, templated insertions generated by prime editing, locus-specific repair biases, long-read sequencing noise, and amplification artifacts collectively challenge conventional amplicon-based analysis workflows.

The analytical landscape for genome editing is currently fragmented. Existing command-line tools, while powerful, often require specialized bioinformatics expertise, creating a bottleneck for many laboratories (*Clement et al., 2019; Amit et al., 2021; Tommaso et al., 2017*). More accessible web-based tools, particularly those running on a server, lack scalability and might raise concerns about data privacy (*Clement et al., 2019, Park et al., 2017*). While other browser-native or locally implemented tools offer improved data protection, they are frequently optimized only for short-read Illumina genotyping, with either absent (*Schmid-Burgk et al., 2014*) or only partial adaptation (Nguyen et al., 2022) to the distinct error profiles of long-read platforms like ONT sequencing.

In this study, we present CleanFinder as a modular and scalable analytical framework designed to address this diversity through anchor-guided localization, constrained semi-global alignment, dual-reference scoring for complex edits, integrated repair pathway annotation, and allele-aware quality control.

In our previous attempt (Nguyen et al., 2022) and during the early development stages of CleanFinder, edit reconstruction relied primarily on adaptive local alignment centered on the anticipated editing site. While local alignment (*Smith–Waterman, 1981*) provides sensitive detection of high-scoring edit-containing subsequences, its design inherently reports only the best local match and may therefore truncate flanking sequence context. In the setting of complex editing outcomes, such as large insertions, composite indel–substitution events, or long-read sequencing data, this behavior required additional post-alignment layers to reconstruct full-length edit structure and ensure accurate classification. Furthermore, classic local alignment is computationally intensive. Conversely, conventional global (Needleman–Wunsch) alignment enforces end-to-end matching of both sequences and does not naturally accommodate terminal overhangs, limiting its suitability for long, error-prone ONT reads where adapters and large indels are common (Needleman et al., 1970) (**Table S2**).

To overcome the structural and computational limitations of the earlier buffer-based approach, we adopted a constrained semi-global (“glocal”) alignment strategy (Durbin et al., 1998). In this framework, short, high-confidence anchor sequences first localize each read to a candidate reference window. Semi-global alignment is then performed within this window without penalizing terminal gaps, preserving full-length read continuity while tolerating variable read boundaries and large indels. By restricting alignment to biologically relevant regions, this anchor-guided design reduces unnecessary matrix evaluation and eliminates the need for manually defined buffer zones. Crucially, the constrained alignment focuses on the biologically relevant segment without discarding reads based on unaligned terminal sequence, ensuring the read remains available for visualization and manual review even in cases of complex or ambiguous editing outcomes. The advantages of this architecture are particularly evident in long-read datasets. ONT reads frequently contain adapter remnants, variable overhangs, and length asymmetries relative to the reference amplicon. For instance, in mtDNA editing experiments, where amplicons reached 2–3 kb, anchor-guided semi-global alignment enabled direct processing of long reads without trimming or window redefinition, while maintaining accurate structural reconstruction. Furthermore, this approach demonstrated high concordance with the dedicated analysis pipeline Mitopore for heteroplasmy quantification (Dobner et al., 2024). The platform was further validated using PacBio HiFi long-read data, where it successfully identified known single-nucleotide polymorphisms (SNPs) within target loci (data not shown).

To further improve scalability, CleanFinder incorporates a k-mer–based pre-filter that enriches for on-target reads and restricts full semi-global alignment to sequences containing locus-specific seeds (*Durbin et al., 2009, Langmead et al., 2012, Li et al., 2018*). This substantially reduces the effective alignment search space and improves throughput across both short- and long-read datasets. Together, anchor localization and k-mer filtering allow CleanFinder to balance alignment fidelity with computational efficiency, enabling analysis of clonal validation experiments as well as multiplexed screening datasets.

The increasing complexity of genome editing modalities, like Prime Editing (PE), introduced further analytical complexity. For instance, PE can generate substitution-only edits, precise insertions, deletions, and composite architectures within the same locus (*Anzalone et al., 2019, Choi et al., 2021, Antoniou et al., 2024, Chauhan et al., 2025*). At the same time, alignment against a single reference sequence is often insufficient. To address this, CleanFinder implements a dual-reference strategy in which each read is aligned independently against both the wild-type reference and the intended edited reference, followed by competitive scoring. The relative alignment scores are then used to assign reads to wild-type, intended edit, or alternative outcomes. This competitive scoring framework improves the classification of complex edits and prime editing products, reliably resolving outcomes such as Reverse Transcriptase Template (RTT)-derived insertions combined with deletions and partially integrated alleles, without requiring manual curation.

Beyond structural classification, CleanFinder integrates mechanistic annotation through RIMA-based repairome analysis (*Taheri-Ghahfarokhi et al., 2018; López de Alba et al., 2025*). Microhomology length, deletion architecture, and templated insertion signatures are extracted directly from aligned alleles, enabling inference of classical non-homologous end joining, microhomology-mediated end joining, and related repair processes without additional experimental input. Embedding this pathway-level annotation within the primary analysis pipeline extends genotyping beyond frequency measurement toward mechanistic interpretation, which is critical for facilitating comparative analyses and for high-impact applications, such as identifying compounds that modulate specific DNA repair pathways during screening.

To demonstrate the framework’s utility in systematic perturbation studies, we conducted a proof-of-concept primary small-molecule screen to identify modulators of prime editing efficiency across substitution- and insertion-based designs, which represent mechanistically distinct repair outcomes (Figure 5**; Figure S5**). In fact, reverse transcription template (RTT)–mediated insertions require successful flap resolution and integration of extended sequence tracts, whereas base substitutions involve shorter sequence replacement events that may engage partially overlapping but distinct repair intermediates (*Anzalone et al., 2019; Peterka et al., 2022*). The observed enhancement was generally marginal, reflecting the highly optimized PEmax system and ePEG RNAs (Nelson et al., 2022). The screen’s biological specificity was confirmed by the strong suppressive effect of known reverse transcriptase inhibitors such as Rilpivirine and Tenofovir alafenamide, which reduced prime editing efficiency by more than 70%, consistent with the reverse transcription–dependent mechanism of PE. Although prior reports have described context-dependent modulation by histone deacetylase (HDAC) inhibitors, characterized by increased insertion efficiencies but decreased substitution efficiencies (Liu et al., 2022), our results revealed a different pattern. Specifically, treatment with HDAC inhibitors such as Nexturastat A, Abexinostat, and Vorinostat led to a consistent reduction in editing efficiency across both substitution and insertion designs (Figure 5 **and Figure S5**). To address this discrepancy, we re-tested selected compounds using the original *HEK3* pegRNA from the primary screen as well as an independent pegRNA targeting the *VHL* locus. Across both loci, HDAC inhibition reproducibly resulted in moderate dose-dependent suppression of prime editing, with sigmoidal response curves and low micromolar ED50 values (**Figure S6)**.

These findings suggest that the impact of chromatin-modifying agents on prime editing may be strongly dependent on locus architecture, chromatin context, pegRNA design, and cellular background. Rather than indicating a universal enhancement or suppression mechanism, our data support a model in which HDAC inhibition modulates accessibility or repair pathway engagement in a context-specific manner. Importantly, the reproducibility of graded responses across independent targets and cell types underscores the robustness of CleanFinder in detecting subtle but biologically meaningful modulation of editing efficiency at screening scale.Conversely, Nimesulide and ROCK inhibitors (Y-27632, CCT137690) increased editing. The effect of ROCK inhibitors is potentially attributable to enhanced cell survival rather than direct editing modulation (Watanabe et al., 2007). DNA-PK inhibition (e.g., AZD7648) did not enhance editing efficiency in our system. Unlike the DSB-generating prime editors described by Dacquay et al. (2025), our nickase-based editor (pCMV-PEmax) does not induce substantial DSBs, likely accounting for the different results.

In addition to edit classification, CleanFinder incorporates allele-aware dropout detection using heterozygous SNP tracking. Whereas most amplicon analysis pipelines focus on indel frequency and deletion size distributions, they do not assess whether such events segregate with loss of one parental allele. Large deletions or primer-site disruptions can result in preferential amplification of a single allele, leading to apparent monoallelic genotypes (*Kosicki et al., 2017; Simkin et al., 2022*). By phasing heterozygous SNPs with deletions on long-read sequencing data (e.g., ONT, PacBio), CleanFinder leverages endogenous polymorphisms as genomic barcodes to distinguish true monoallelic editing from amplification bias or loss of heterozygosity. These informative SNPs can be identified a priori using the WSABI platform (Dobner et al., 2026, in preparation) or other available tools, facilitating selection of high-confidence heterozygous markers in defined cell-line backgrounds. This allele-aware framework provides an additional quality-control layer particularly valuable for long-read sequencing and clonal validation workflows.

CleanFinder is available in two complementary environments sharing the same full-featured analytical engine: a Python-based command-line interface (CLI) and a high-resolution mode within the browser-native application. Both provide semi-global alignment, dual-reference scoring, batch processing, configurable editing modes, and downstream annotations. The modular, scriptable CLI is designed for community extension and integration into high-throughput workflows, supports large files beyond browser memory limits, and enables structured CSV exports for downstream automation.

In contrast, the browser-native Turbo module implements heuristic anchor-guided localization to enable rapid genotype categorization and interactive visualization (*Lipták et al., 2022*). By prioritizing bounded-mismatch anchor scanning and restricting dynamic programming to selected contexts, Turbo achieves substantially higher processing speeds suitable for exploratory, plate-scale analysis. This acceleration comes with a deliberate reduction in downstream annotation: Turbo focuses on rapid categorization and visualization, whereas the CLI provides comprehensive alignment-dependent summaries and mechanistic metrics. The coexistence of these two implementations reflects differing analytical needs, from rapid screening to in-depth mechanistic interrogation.

### Limitations and future directions

Several limitations should be acknowledged. The small-molecule screen was conducted at a single concentration (10 µM) to demonstrate analytical scalability under high-throughput conditions. Although single-dose primary formats are widely used for initial hit triage in screening campaigns (Butkiewicz et al., 2017), this approach does not establish compound potency, specificity, or fully resolve potential cytotoxic effects. While overtly cytotoxic compounds were identified and excluded based on phenotypic assessment, comprehensive viability profiling was beyond the scope of the primary screen. Secondary validation, including dose–response analysis and orthogonal assays, is therefore required to confirm reproducibility, disentangle toxicity from true editing modulation, and establish mechanistic relevance of the identified modulators.

Across both Illumina and ONT datasets, cross-platform benchmarking demonstrated strong concordance of editing outcomes. However, ONT sequencing remains subject to intrinsic platform-specific basecalling biases, particularly within homopolymeric regions where indel rates are artificially elevated. These effects reflect characteristics of the sequencing technology rather than the analytical framework itself. Future CleanFinder versions will incorporate optional warning flags for deletions occurring within homopolymeric stretches, especially when overlapping predicted editing windows, to aid interpretation in long-read datasets. Until such safeguards are implemented, results derived from homopolymeric regions should be interpreted with appropriate caution. Recent advances in sequencing technologies further enhance amplicon-based genome editing analyses. Updated short-read platforms such as the Illumina MiSeq i100 improve performance for low-diversity CRISPR libraries, while compact long-read devices and low-cost flow cells such as ONT’s Flongle make targeted sequencing accessible to smaller laboratories. Frameworks such as CleanFinder that support both high-accuracy short reads and cost-efficient long-read datasets are therefore increasingly important for flexible and accessible genome editing analysis.

The anchor-guided semi-global alignment strategy preserves full-length read continuity and enables robust structural reconstruction. In rare cases, minor labeling inconsistencies may occur in the raw output string representation of complex alleles; however, the full alignment context is retained, allowing manual confirmation of structural integrity when needed.

CleanFinder was intentionally developed as a modular, open-source framework to facilitate ongoing extension and community-driven development. Planned enhancements further include support for multi-gRNA analysis within the browser-native implementation and expanded reporting functionality within the command-line interface, including automated generation of publication-quality graphical summaries.

Future browser-native versions may incorporate client-side parallelization (e.g., Web Workers) to enable concurrent processing of multiple input files across available CPU cores without altering the deterministic alignment and classification logic. Given the modular pipeline architecture, such parallelization can be implemented without affecting reproducibility or interpretability.

Taken together, CleanFinder represents a methodological progression from buffer-based local alignment toward an anchor-guided, semi-global, and modular architecture capable of resolving diverse genome editing outcomes across sequencing platforms and experimental scales. By integrating structural reconstruction, competitive dual-reference scoring for complex edits, k-mer–guided acceleration, repair pathway annotation, allele-aware quality control, and scalable deployment modes, the framework provides a flexible and extensible foundation for contemporary genome engineering workflows.

## Material and methods

### Cell Culture

HEK293T and HaCaT cells were cultured in Dulbecco’s Modified Eagle Medium (DMEM; Gibco) supplemented with 10% fetal bovine serum (FBS; Gibco) and 1% penicillin-streptomycin.

Human erythroleukemia K562 cells were cultered in RPMI 1640 containing glutamine (Gibco) supplemented with 10% fetal bovine serum (Gibco), 2% Hepes (Life Technologies), 1% sodium pyruvate (Life Technologies), and 1% penicillin and streptomycin (Life Technologies) at 37°C and 5% CO2.

Human male KOLF2.1J (Jackson Laboratory), IMR90 (WiCell) and female iPS12 (Cell Applications) iPSCs were cultured on 1% Geltrex-coated plates (Gibco) in mTeSR™ Plus medium (STEMCELL Technologies) supplemented with 5X Supplement and 1% penicillin-streptomycin (complete medium), at 37°C in a humidified atmosphere containing 5% COLJ. For routine maintenance, iPSCs were passaged using enzyme-free ReLeSR (STEMCELL Technologies). To generate single-cell suspensions, cells were dissociated with Accutase (Pan Biotech) and replated in complete medium supplemented with the CEPT cocktail (R&D Technologies).

All cells used in this study were tested for Mycoplasma contamination using the Mycoplasma Detection Kit (SouthernBiotech), following the manufacturer’s instructions.

The human male iPSC line BIHi-005-A-24 (Lisowski et al., 2024) and its derivatives were routinely monitored for mycoplasma contamination by PCR using nine primers to amplify the six most common mycoplasma strains. The positive control and internal control were kindly provided by Dr. Cord Uphoff (DZMS, Germany). The cells were cultured on Geltrex (Gibco)-coated plates using StemMACS iPS-Brew XF medium (Miltenyi Biotec), supplemented with MycoZap (Lonza) in a humidified atmosphere of 5 % CO2 at 37 °C and 5 % oxygen. They were passaged at 70-80 % confluence with 0.5 µM EDTA (Invitrogen) in 1 x PBS (Gibco) and replated in complete medium supplemented with the CEPT cocktail (R&D Technologies).

Quality control of iPSCs was performed as previously described (Dobner et al., 2024). Briefly, Karyotyping was performed at the Institute of Human Genetics and Anthropology, Heinrich Heine University. Low-resolution karyotyping was conducted using the Illumina Infinium Global Screening Array (GSAMD-24v3-0-EA_20034606) by Life&Brain. Pluripotency was assessed using hiPSCore (Dobner et al., 2024).

Human non-mobilized peripheral blood CD34LJ HSCs were obtained from patients with sickle cell disease (Necker–Enfants malades Hospital, Paris, France) following written informed consent and in accordance with the Declaration of Helsinki (IRB approval: DC-2024-6899, CPP Ile-de-France II). CD34LJ cells were purified using the CD34 MicroBead Kit (Miltenyi Biotec), thawed, and cultured for 24h at 5 × 10LJ cells/ml in StemSpan medium supplemented with penicillin/streptomycin, 250 nM StemRegenin1, and recombinant human SCF (300 ng/ml), Flt-3L (300 ng/ml), TPO (100 ng/ml), and IL-3 (60 ng/ml).

For editing, 2 × 10LJ cells per condition were electroporated with 6.4 μg PEnmax mRNA and 200 pmol synthetic pegRNA (IDT) using the P3 Primary Cell 4D-Nucleofector X Kit S (Lonza; program CA-137). Cells were recovered for 10 min at 37°C and subsequently cultured for 6 days prior to DNA extraction. Tris-EDTA–transfected cells served as negative controls.

### Genome Engineering

The pSpCas9(BB)-2A-GFP (PX458) plasmid was obtained from Addgene (#48138). Guide RNAs (gRNAs) for Cas9 and Cas12 were designed using the CHOPCHOP (Labun et al., 2019) online tool (https://chopchop.cbu.uib.no/). Cas9 gRNAs were cloned into the PX458 vector. Cas9 gRNA sequences are listed in the **Table S3** For knock-in experiments, donor templates containing the desired point mutations were designed and synthesized as single-stranded DNA oligonucleotides by Integrated DNA Technologies (IDT). The sequences of donor oligos are also provided in **Table S3**. For prime editing, the pCMV-PEmax plasmid was obtained from Addgene (#174820). To generate a GFP-tagged version, the EGFP gene was PCR-amplified from PX458 and cloned into AgeI-linearized pCMV-PEmax using the CloneExpress II kit (Vazyme). Prime editing guide RNAs (pegRNAs) targeting *EMX1*, *XPA*, *VHL*, *POU5F1* and *HEK3 (***Table S3**) were cloned into the pU6-pegRNA-GG-acceptor vector (Addgene #132777) or the pU6-tevopreq1-GG-acceptor vector (Addgene #174038). All constructs were verified by Sanger sequencing. Genome engineering experiments were performed as previously described, with minor modifications (Ramachandran et al., 2021).

To generate gene knockouts in iPSCs, guide RNAs were transfected using Lipofectamine Stem Reagent (Thermo) according to the manufacturer’s instructions. For homology-directed repair (HDR)-mediated knock-ins, guide RNAs were co-transfected with single-stranded donor oligonucleotides under the same conditions. For prime editing, pegRNAs were co-transfected with the pCMV-PEmax plasmid. In HEK293T cells, genome editing reagents were delivered using FuGENE® HD Transfection Reagent (Promega), following the manufacturer’s protocol. HaCaT cells were transfected using the Neon™ Electroporation System (Thermo Fisher Scientific) at 1650V with a 10ms pulse width and 1 pulse. K562 cells (106 cells/condition) were transfected with 3.6 μg of PEmax or PEnmax and 1.2 μg of the pegRNA-containing plasmid using AMAXA Cell Line Nucleofector Kit V (VCA- 1003, Lonza) and U-16 program (Nucleofector 2b, Lonza). For CRISPR-Cas12a (Cpf1) mediated genome editing experiments, the Cas12a protein, guide RNA, and donor oligonucleotides (all purchased from IDT) were assembled into ribonucleoprotein (RNP) complexes and delivered via electroporation using a Neon Electroporation system (Thermo Scientific). Electroporation was performed at 1400V, with a 20ms pulse width and 1 pulse. Sequences of Cpf1 guide RNAs and donor oligos are listed in **Table S3**. At 48 hours post-transfection (excluding Cpf1 electroporations) GFP-positive cells were isolated by fluorescence-activated cell sorting (FACS). Bulk-sorted cells were expanded and plated at limiting dilution into 96-well plates to derive clonal populations. Duplicate plates were prepared: one for continued culture and one for genomic DNA extraction. DNA extraction was performed using Proteinase K. The enzyme was added to the reaction at a final concentration of 50 µg/mL and incubated at 50 °C for at least 1 hour. Proteinase K was then inactivated by heating the reaction to 95 °C for 5 minutes. 20 ng of genomic DNA (gDNA) were used for PCR amplification and subsequent genotyping analysis. For prime editing and base editing of the *HBG1/2* promoters, genomic DNA was isolated from CD34LJ HSCs 6 days post-transfection using the PureLink Genomic DNA Mini Kit. The HBG1/2 promoter regions were amplified with Phusion Flash High-Fidelity Mastermix in 15 μL reactions containing 1.5 μL DNA and 0.2 μM barcoded primers. PCR conditions were 98 °C for 3 min; 30 cycles of 98 °C for 10 s, 60 °C for 20 s, and 72 °C for 30 s; followed by 72 °C for 2 min. Amplicons were purified (HighPrep PCR Clean-up System) and assessed by fragment analyzer. Illumina indexes were added in a second PCR using KAPA HiFi HotStart Ready Mix with 0.067 ng template and 0.5 μM indexed primers (25 μL reaction). Cycling conditions were 72 °C for 3 min; 98 °C for 30 s; 10 cycles of 98 °C for 10 s, 63 °C for 30 s, and 72 °C for 3 min; and 72 °C for 5 min. After purification and quality control, libraries were quantified by Qubit, sequenced on an Illumina NextSeq, demultiplexed with bcl2fastq, and analyzed using CleanFinder. For generation of the DdCBE plasmids required for mtDNA editing, TALE sequences were designed using TALEN Targeter (https://tale-nt.cac.cornell.edu/node/add/talen) and Paired Target Finder (https://tale-nt.cac.cornell.edu/node/add/talef-off-paired). The TALE sequences were ordered as gene synthesis from GeneArt (Thermo Fisher Scientific) and codon optimized for human hosts. They were cloned into Addgene plasmids #158096 and #158092 via golden gate cloning using Esp3I (NEB) and fluorescence marker (mCherry, Neongreen) were added to the end of the constructs linked via T2A sequences. The plasmids were verified via Sanger sequencing.

BIHi-005-A-24 iPS cells were co-transfected with both DdCBE plasmids using ScreenFect A-Plus according to manufacturer’s instructions. At 48 hours post-transfection, cells positive for both Neongreen and mCherry were sorted by FACS and seeded in low density (300 and 500 cells) to allow for growth of single colonies. Approximately 10-12 days after seeding, single colonies were picked and partly used for further cultivation, the other part used for an initial screen via Sanger sequencing (Eurofins). For Sanger sequencing analysis, the cells were lysed using Phire Animal Tissue Direct PCR Kit (Thermo Fisher Scientific) following the dilution protocol. Lysates were used as template in a PCR reaction to amplify the target region with Q5 High-Fidelity DNA Polymerase (NEB) resulting in amplicons of about 1kb size.

A complete list of oligonucleotides is provided in **Table S4**.

### Prime Editing Screening

Small molecules from Selleck and Sigma libraries were obtained as 10 mM DMSO stocks. Primary working stocks were prepared at 100 µM in DMEM lacking serum and antibiotics, following the manufacturers’ guidelines. Working plates were stored at −20 °C for up to three months. To assess prime editing efficiency, HEK293T cells were reverse-transfected with pegRNA and pCMV-PEmax plasmids. Several transfection reagents, including PEG-based systems, were evaluated; FuGENE (Promega) was selected for its high transfection efficiency and low cytotoxicity in HEK293T cells. The DNA-FuGENE complex was incubated with the cell suspension for 5LJminutes before plating into three replicate 96-well plates pre-coated with poly-L-ornithine to promote adherence and prevent detachment during medium changes. All washing steps were performed using PBS containing calcium and magnesium to preserve cell integrity and adhesion. Immediately after plating, cells were treated with small molecules at a final concentration of 10 µM or with DMSO as a vehicle control. Plates were incubated for ∼48 hours at 37 °C in a humidified atmosphere with 5% CO_₂_.

Cell health and transfection efficiency were monitored using a Leica DMi1 inverted microscope. EGFP expression was used to assess transfection rates, and cellular morphology and confluency were visually evaluated to identify signs of cytotoxicity. To assess potential cytotoxicity under our experimental conditions, we quantified cell numbers by counting nuclei in compound-treated samples compared to DMSO controls using Fiji (Image J) software as previously shown (*K*ř*ížkovská et al., 2023*).

Genomic DNA was extracted 48 hours post-treatment, and prime editing efficiencies were quantified by next-generation sequencing using the pipeline.

Following analysis with CleanFinder, output files (CSV, Excel format and HTML) were generated containing the proportion of edited versus unedited (wild-type, WT) reads for each treatment condition, as detailed in the supplementary data. For each experimental condition, three replicates were analyzed, and normalized to editing percentages of the DMSO control on the same plate. Compounds that increased editing efficiency to >120% relative to the DMSO baseline were classified as enhancers, whereas compounds that reduced efficiency to below 70% were classified as inhibitors. To accommodate for data variability, robust z-scores were calculated using the median and median-absolute-deviation (Rindskopf et al., 2010), and meaningful effects were inferred for robust z-scores ≥ 2. A complete list of hits from the drug screening is provided in **Table S5.**

### Library preparation and deep sequencing

Illumina DNA amplification was performed as previously described (*Ramachandran et al. 2021*). Briefly, first-level PCR reactions were performed using 1 μl of lysate as a template in a 6.25 μl Phanta (Vazyme) PCR reaction according to the manufacturer’s protocol (annealing temperature: 60°C; elongation time: 15 s, 19 cycles). PCR custom primers were designed to have adaptors for the second PCR. From this reaction, 2 μl were transferred to a second-level PCR using the same cycling conditions and a barcoded primer unique for each clone.

PCR products were pooled, size-separated using a 1% agarose gel and purified using the GeneJET gel extraction kit (Thermo Fisher Scientific). Illumina libraries were sequenced on an Illumina MiSeq benchtop sequencer using a Nano v2 reagent kit in a single-end, 251-cycle run. Libraries were loaded at 6 pM with a 10% PhiX spike-in to balance low-diversity libraries. Additional sequencing was performed on a MiSeq i100 Plus system using patterned flow cells (5 million, 300-cycle kit). A total of 251 single-end cycles were sequenced. Libraries were loaded at 60 pM. Base calling and BCL-to-FASTQ conversion were performed onboard using BCL Convert (v3.6.3 and v4.0.3).

ONT DNA amplification was performed as previously described (*Nguyen et. al., 2022*).

The ONT sequencing library was prepared using the Ligation Sequencing Kit (SQL-LSK109) or the SQK-NBD114.24 Sequencing Kit (for barcoding) according to the manufacturer’s protocol, loaded at 10 fmol and sequenced using a MinION MK1C or MinION using a Flongle flow cell and adapter.

All sequencing libraries were quantified using Qubit4 (Thermo) and Tapestation 4150 (Agilent). Illumina libraries were basecalled and demultiplexed onboard the instrument, and the resulting files were analyzed using CleanFinder.

Raw ONT POD5 data was basecalled using dorado (0.4.3+656766b) using the super accuracy model (dna_r10.4.1_e8.2_400bps_sup@v4.2.0). For ONT sequencing of CRISPR/Cas genome editing experiments, mean read depth was 75,756 (50,534 - 120,458), mean read length was 534.6 (443.9 - 616.3), and mean PHRED per read was 17.5 (14.9 - 26.5). For ONT sequencing of mtDNA BE experiments, mean read depth was 1,128 (630 - 1,548), mean read length was 5,579,7 (4,369.4 - 7,062.4), and mean PHRED per read was 20.5 (19.3 - 21.7).

Necessary sequencing depth depends on several factors which need to be considered: Detection probability depends on allele frequency and read depth according to a binomial model (probability of detection = 1 − (1 − p)^N). For example, reliable detection (∼95% probability) of a 1% allele requires ∼300 reads, whereas detection of a 0.1% allele requires ∼3,000 reads; single-read events (e.g., 0.02% in 5,000 reads) may be retained but should be considered indicative rather than statistically robust.

### CleanFinder implementation and Alignment algorithm

CleanFinder is implemented as a browser-native single-page application (SPA) using standard web technologies (HTML5, CSS3, and JavaScript). In browser-native modes (standard and Turbo), all sequence processing, including anchor detection, constrained semi-global alignment, genotype classification, and visualization, is executed entirely client-side within the user’s web browser. This architecture preserves data privacy and eliminates the need for server-side computation or dedicated software installation. When gene-based analysis is selected, gene symbols are resolved to genomic coordinates using the Ensembl REST API, and corresponding reference sequences are retrieved programmatically from Ensembl. Alternatively, users may provide custom reference sequences directly. Interactive visualizations are rendered using Chart.js, while gene schematics and alignment views are generated as Scalable Vector Graphics (SVG). Input data consist of FASTQ files generated from targeted amplicon sequencing experiments (Illumina, PacBio, or ONT). Reads are processed iteratively to enable analysis of large datasets within browser environments.

Read placement is determined using short reference-derived anchor sequences located at defined positions within the amplicon. Both forward and reverse-complement anchors are evaluated to determine read orientation. Reads lacking valid anchor matches within the allowed mismatch threshold are excluded from downstream alignment.

Following anchor localization, reads are aligned using a constrained semi-global (glocal) dynamic programming strategy. Alignment is restricted to the anchor-defined reference window, and terminal gaps are not penalized. This approach preserves full-length structural continuity while tolerating variable read boundaries and large insertions or deletions. By limiting alignment to biologically relevant regions, computational complexity is reduced relative to full-length global alignment. For prime editing experiments, reads are independently aligned against both the wild-type reference sequence and the intended edited reference sequence using the same constrained semi-global strategy. Comparative alignment scoring is used to assign reads to wild-type, intended edit, or alternative categories. This dual-reference framework improves classification of complex architectures, including composite insertion–deletion events and partially integrated reverse transcription template products.

To improve scalability, CleanFinder optionally applies a locus-specific k-mer prefilter prior to dynamic programming alignment. Reads that do not contain predefined seed sequences (10 bases for Illumina and 13 for ONT) derived from the reference are excluded from alignment, thereby reducing computational load for large datasets.

The Turbo mode is a browser-native configuration optimized for rapid plate-scale screening. In Turbo mode, genotype categorization relies primarily on anchor detection and rule-based evaluation of internal sequence segments rather than full dynamic programming alignment for every read. This heuristic approach increases throughput while reporting a reduced set of alignment-derived annotations compared to the standard workflow.

The performance of Turbo mode was evaluated across several hundred experimental and simulated datasets encompassing CRISPR/Cas KO, knock-in KI, BE, and PE experiments. Across simple indel-based editing scenarios, particularly typical Cas-mediated KO experiments characterized by single deletions near the cut site, Turbo mode demonstrated high concordance with the full CleanFinder alignment pipeline.

However, due to its rule-based classification strategy, Turbo mode may be less suited for complex editing architectures involving combinations of multiple events within the same read (e.g., concurrent deletions, insertions, and substitutions, or large templated insertions). In such cases, the standard alignment-based workflow provides more granular annotation and improved resolution of compound alleles. Consistent with its design goal of rapid screening, Turbo mode provides a streamlined output focused on edit categorization and frequency estimation, whereas the full workflow generates comprehensive alignment metrics, positional variant mapping, and extended annotation layers. Example test datasets are publicly available at https://cleanfinder.org.

The Python CLI implements the full alignment and classification workflow and supports batch processing of large datasets. It exposes parameters for anchor length, mismatch tolerance, and alignment mode, and produces structured output files suitable for integration into downstream analytical pipelines.

A comprehensive user manual (CleanFinder 1.0 Manual) describing configuration options, analysis modes, and output interpretation is available within the web application (“Info” section) and via the project repository on GitHub (https://github.com/andrearossi-lab/CleanFinder/tree/main) and on cleanfinder.org.

### Allelic Dropout

To detect potential hemizygous loss in clonal cell lines, we implemented an Allelic Dropout SNP Analyzer module. The analysis requires three primary inputs: long-read sequencing data (FASTQ), a wild-type reference amplicon sequence (optionally retrieved via the gDNA Finder integrated module), and a user-defined stable anchor sequence (∼20 bp). This anchor serves as a conserved genomic landmark, ideally situated distal to the expected editing site, to enable orientation-independent read filtering. The computational pipeline executes a multi-stage workflow: (1) Read Filtering: Raw FASTQ data are scanned for the anchor sequence and its reverse complement to isolate on-target reads. (2) Matrix Construction: A user-defined subset of anchor-positive reads undergoes affine-gap alignment against the reference to construct a positional base-frequency pileup (matrix). (3) SNP Interrogation: The algorithm scans this matrix for heterozygous candidates, defined by a stoichiometric balance (default frequency range: 30–70%) of two dominant alleles and a minimum coverage threshold. SNP candidates are ranked using a composite quality score that prioritizes positions with high read-depth and allele frequencies approaching 50% (maximal symmetry). This scoring penalizes low-coverage sites and highly skewed allele distributions, which are more likely to represent sequencing noise rather than true heterozygous variants. The highest-confidence SNP is reported with a comprehensive visual profile including its genomic coordinates, biallelic identities, stoichiometric ratios, and a 21-bp flanking context (10 bp up/downstream) for sequence-level verification. Example test datasets are publicly available at https://cleanfinder.org (Allelic Drop folder).

### Genotype Quantification, Repair Annotation, and Output

Following constrained semi-global alignment and genotype assignment, reads are grouped by identical allele structures and quantified to generate genotype frequency distributions. User-defined frequency thresholds can be applied to filter low-frequency variants. Deletion and insertion events are extracted directly from aligned allele structures. For indel-based editing, repair-associated features, including microhomology length, insertion duplication, and deletion boundaries relative to the PAM site, are computed to enable RIMA-based repairome classification. Deletions are annotated as PAM-proximal or PAM-distal based on their position relative to the predicted cleavage site, facilitating mechanistic interpretation of repair patterns. In the browser-native implementation, results are rendered as interactive visualizations, including allele tables, frequency plots, and summary charts. These outputs can be downloaded as publication-ready graphics and exported in structured tabular formats (CSV and Excel) for downstream analysis. The full session state, including input parameters and computed results, can be stored as a self-contained HTML file for reproducibility.

The Python CLI implements the same analytical workflow and prints structured output to the console while generating tabular result files (including Excel-compatible formats) suitable for batch processing and integration into external pipelines.

### Correlation Analysis

To assess the agreement between ONT and Illumina platforms, correlation analyses were performed on percentages of the top 10 shared genotypes across individual clones (55 different genotypes) and bulk genome-edited samples (45 different genotypes). These experiments spanned 9 different genes across ONT and Illumina sequencing including base substitutions (7 different genotypes), insertions (9 different genotypes), knock-ins (16 different genotypes), deletions (59 different genotypes), and wildtype sequences (9 different genotypes). Pearson correlation coefficients were calculated using the open-source statistical software R (version 4.5.2) using the package ggpubr (version 0.6.2, alternative hypothesis: “two-sided”), and visualized using the R package smplot2 (version 0.2.5).

### Integration of Rational Indel Meta-Analysis 2 (RIMA2) and REPAIRome

CleanFinder integrates repair pathway inference directly into the genotype reconstruction workflow (*Taheri-Ghahfarokhi et al., 2018, EL de Alba · 2025 et al., 2025).* Repair annotation is computed from aligned allele structures generated by the constrained semi-global alignment, without requiring external preprocessing or post hoc file conversion. For each aligned allele containing a deletion event, the software identifies the deletion junction within the alignment string and extracts the deleted reference sequence. Microhomology (MH) is then evaluated by comparing the deleted sequence to flanking reference segments immediately upstream and downstream of the deletion boundary. The maximal contiguous sequence identity at the junction is recorded as the MH length. Deletions with microhomology length ≥2 bp are classified as microhomology-mediated end joining (MMEJ), whereas deletions with no detectable microhomology (MH = 0) are classified as non-homologous end joining (NHEJ). Single-base deletions are treated as NHEJ events. Deletion events are additionally annotated according to their positional relationship to the predicted cleavage site. Based on the distance between the deletion boundary and the cut site, deletions are categorized as PAM-proximal (≤5 bp from the cut site) or PAM-distal (>5 bp). This positional stratification enables inference of locus-dependent repair signatures and cleavage architecture effects. Microhomology lengths are aggregated across all reads to generate a microhomology length distribution. In parallel, repair pathway proportions (NHEJ vs MMEJ) are computed by summing allele frequencies according to MH classification. These metrics are visualized in the browser-native implementation as interactive bar and pie charts and are exported in structured tabular formats in both the web application and the Python CLI. Insertion events are parsed directly from the aligned allele strings and quantified as part of the genotype distribution. While insertion duplication status and templated characteristics can be inferred from alignment context, primary pathway classification within CleanFinder focuses on deletion-associated microhomology features. This integrated implementation enables repair pathway inference to be computed natively from reconstructed alleles and reported alongside genotype distributions, frame status, and editing efficiency metrics.

## Code and Data Availability

The Python command-line implementation of CleanFinder and representative test datasets used for validation are publicly available. The source code and the Python command-line implementation are accessible via GitHub at: https://github.com/andrearossi-lab/CleanFinder

Test datasets and manual files can be accessed at:

https://iufduesseldorf-my.sharepoint.com/my?id=%2Fpersonal%2Fandrea%5Frossi%5Fiuf%2Dduesseldorf%5Fde%2FDocuments%2FDatasets%5FQReads&ga=1 and https://data.mendeley.com/ (doi: 10.17632/7njkjh73pk.1)

The browser-native version is available at cleanfinder.org

## Computational environment

All analyses were performed using CleanFinder in a local browser environment on an Apple MacBook Pro equipped with an Apple M4 processor (14 cores, 24 GB RAM). Analyses were conducted in Google Chrome (version 145.0.7632.77). Cross-browser compatibility was verified using current versions of Safari, Mozilla Firefox, and Microsoft Edge. Additional analyses were performed on a Lenovo P14s workstation equipped with an Intel Core Ultra 7 165H processor (16 cores, 64 GB RAM) and on a Lenovo ThinkPad laptop equipped with an Intel Core i5-10210U processor (4 cores, 16 GB RAM).

## Acknowledgments

We thank Hanna Spiessbach, Marie Brauers, Katrin Hollweg, Vanessa Baltruschat, and Nadine Teichweyde for their technical support with the experiments. We are grateful to Dagmar Wieczorek for her continuous support.

This project was directly supported by the Deutsche Forschungsgemeinschaft (DFG) (RO5380/12-1 to A.R) and the AFM-Téléthon (25179 to A.M., A.R., and Z.K.).

The Environmental Adaptation and Cellular Resilience lab and the GEMD are also funded by Deutsche Forschungsgemeinschaft (DFG) (RO5380/1-1, RO5380/10-1, RO5380/13-1 and RO5380/14-1 to A.R.), VHL (von Hippel-Lindau) Betroffene Familien e.V. through a grant awarded to A.R, European Union (EU) and North Rhine-Westphalia (NRW) start-up grant (IN-SU-3-001 to A.R. and J.D.), and the Leibniz Competition (SAW) Cooperative Excellence project (K642/2024 to A.R.).

The IUF is funded by the federal and state governments, the Ministry of Culture and Science of North Rhine-Westphalia (MKW), and the Federal Ministry of Education and Research (BMBF).

Figures were generated using Canvas X Draw, Adobe Illustrator and Lucid (Fig. 1) and Biorender (Fig. 2, 3, 4, 5, S1, S2, S3, S4, S5).

## Ethics

The applicant laboratory has the necessary permission to work with recombinant DNA and genetically modified organisms under S1 and S2 conditions.

## Contributions

Conceptualization (A.R.), methodology and development (A.R.), investigation (H.R., S.B., V.I., B.M, T.N., I.T., T.N., J.D.), visualization (J.D., H.R., A.R.), writing - original draft (A.R.), manuscript (A.R. with the input of all authors), statistics (J.D. and C.S.) review and editing (all authors): resources (A.R., Z.K., A.P. and A.M), supervision (A.R., Z.K., A.M., A.P.), project administration, and funding acquisition (A.R., Z.K., J.K., A.P. and A.M.). All authors reviewed and approved the manuscript.

The authors declare no competing interests.

## Supplementary Information

**Table S1.** Feature comparison of CRISPR/Cas editing analysis platforms. Comparison of the CRISPR/Cas editing analysis platforms CleanFinder, OutKnocker, Cas-Analyzer, and CRISPResso2 across key technical specifications, including deployment architecture, sequencing compatibility, and advanced analytical capabilities such as ONT support and specialized pipelines for Base and Prime editing.

| Category | Feature | CleanFinder | OutKnocker | Cas-Analyzer | CRISPResso2 |
| --- | --- | --- | --- | --- | --- |
| Deployment | Platform | Browser-native (local) + CLI | Browser-native (local) | Browser-native (local) | Web (server-side) + CLI |
|  | Local Data Processing | Yes | Yes | Yes | CLI only |
| Throughput | Batch Upload (Web) | 96+ samples | Up to 96 | 1 | 4 (Browser version) |
|  | Batch Processing (CLI) | Yes | No | No | Yes |
| Sequencing | Illumina Support | Yes | Yes | Yes | Yes |
|  | ONT Support | Yes | No | Yes | Yes (requires tuning) |
|  | Speed | Fast | Fast | Fast | Slow (Browser version) |
| Analysis | Handles 100% Editing | Yes | Limited | Yes | Yes |
|  | BE & PE Pipelines | Yes | No | No | Yes |
|  | Delta-size Labeling | Yes | Yes | No | Yes |
|  | Frame Annotation | Yes | Yes | No | Yes |
|  | Repair Pathway Annotation | Yes | No | No | No |
| Advanced | Edit Settings (GAP, etc.) | Yes | No | No (fixed presets) | Yes |
|  | Allelic Dropout | Yes | No | No | No |
|  | Turbo Mode | Yes | No | No | No |
|  | Genome Viewer | Yes | No | No | No |

**Table S2:**
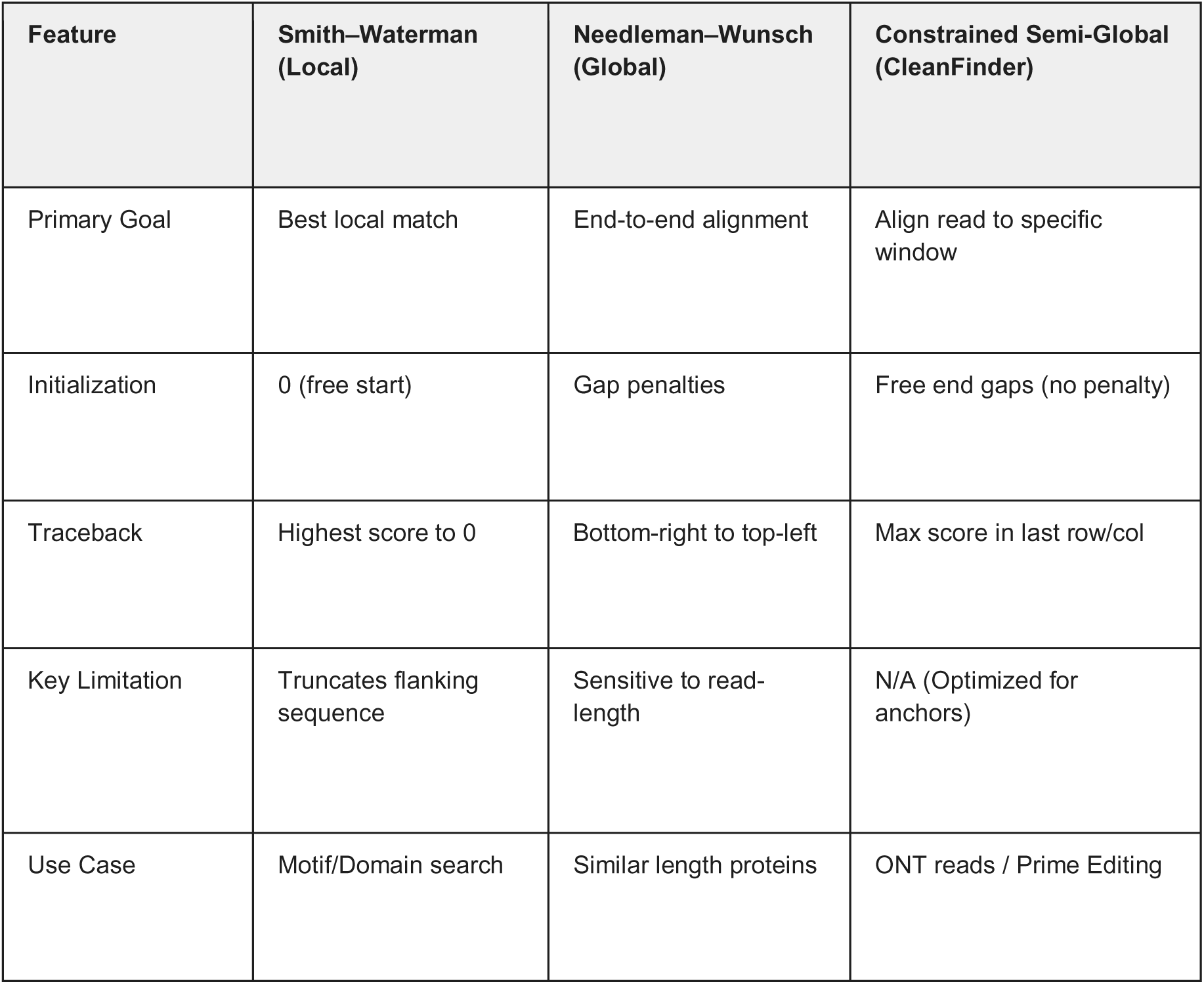
Algorithmic Rationale for Constrained Semi-Global Alignment in CleanFinder. A comparative overview of dynamic programming strategies and their suitability for processing genome editing data.Smith–Waterman (Local): Optimized for identifying conserved motifs; however, the zero-score reset mechanism often truncates flanking context, potentially missing large insertions or deletions at the edges of the editing window. often truncates flanking context, potentially missing large insertions or deletions at the edges of the editing window. Needleman–Wunsch (Global): Requires end-to-end alignment; this becomes computationally prohibitive and introduces alignment artifacts when processing long reads (e.g., ONT) against a specific genomic reference window due to terminal gap penalties. Constrained Semi-Global (CleanFinder): By implementing an anchor-guided localization followed by a semi-global alignment with free end-gaps, CleanFinder ensures that the sequence of interest is aligned entirely within the candidate window. This approach avoids the truncation issues of local alignment and the length-sensitivity of global alignment, providing a robust framework for capturing complex structural variants in Prime Editing and long-read sequencing datasets.

| Feature | Smith–Waterman<br>(Local) | Needleman–Wunsch<br>(Global) | Constrained Semi-Global<br>(CleanFinder) |
| --- | --- | --- | --- |
| Primary Goal | Best local match | End-to-end alignment | Align read to specific window |
| Initialization | 0 (free start) | Gap penalties | Free end gaps (no penalty) |
| Traceback | Highest score to 0 | Bottom-right to top-left | Max score in last row/col |
| Key Limitation | Truncates flanking sequence | Sensitive to read-length | N/A (Optimized for anchors) |
| Use Case | Motif/Domain search | Similar length proteins | ONT reads / Prime Editing |

**Figure S1.**
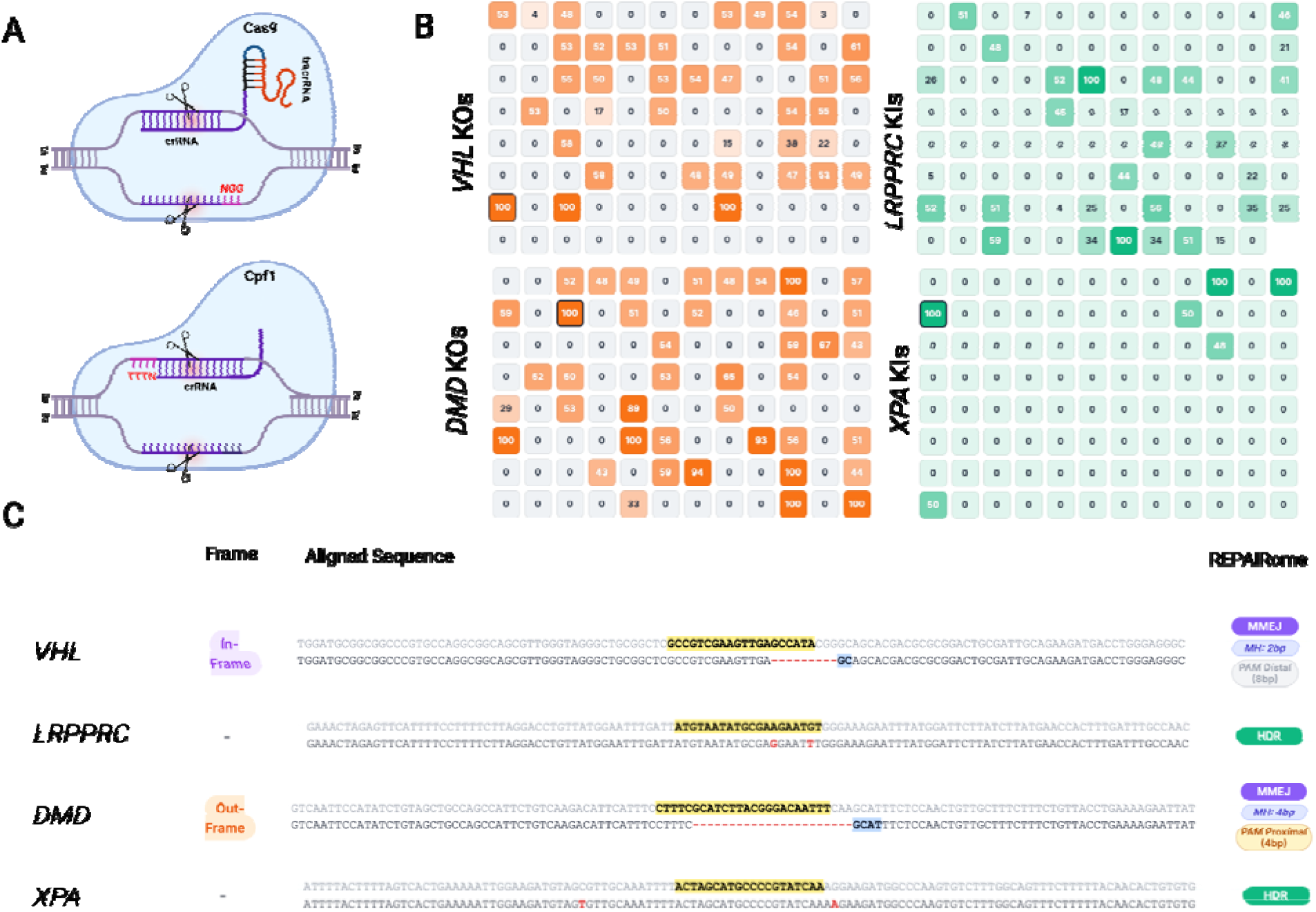
**Cross-validation of CleanFinder across Cas9- and Cas12-mediated knockout and knock-in editing in human iPSCs. (**A) Schematic representation of Cas9 and Cas12 (Cpf1) editing architectures, highlighting differences in guide RNA configuration and PAM orientation. (B) Heatmap visualization of genotype frequencies across representative genome editing experiments in human induced pluripotent stem cells (iPSCs). (C) Cas9- and Cas12-mediated knockout (KO) experiment targeting *VHL* and *DMD*, showing structured annotation of deletion size, frame status (in-frame/out-of-frame), microhomology length (MH), and PAM proximity (proximal/distal), consistent with MMEJ-associated repair. Right panels: Cas9- and Cas12-mediated knock-in (KI) experiments at the *LRPPRC* and *XPA* loci, demonstrating precise homology-directed repair (HDR)–mediated sequence integration. These examples illustrate nuclease- and locus-dependent variation in repair pathway signatures and demonstrate consistent classification of KO and KI outcomes across editing systems.

**Figure S2.**
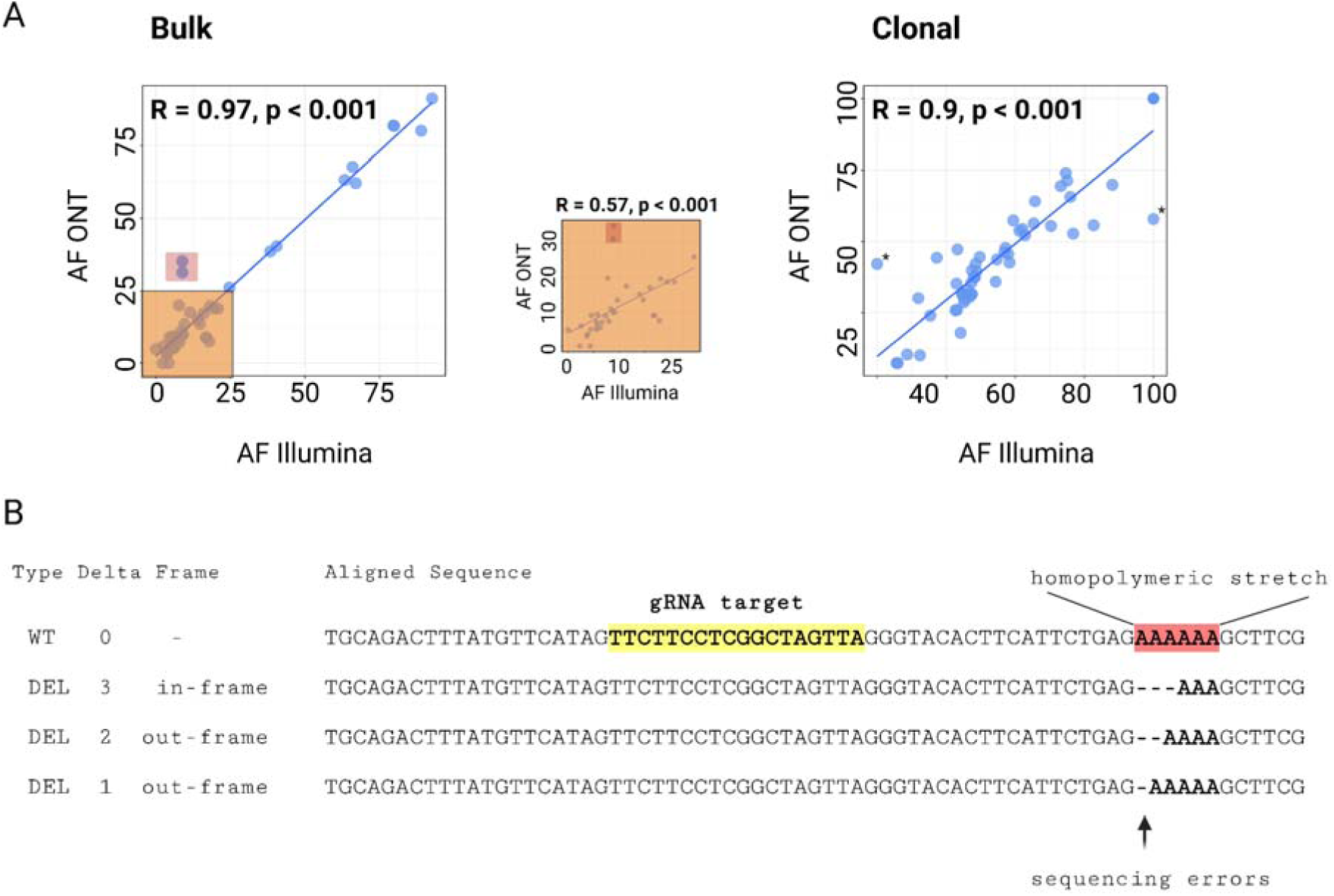
Cross-platform genotype concordance and homopolymer-associated indel artifacts in ONT sequencing. Comparison of allele frequency (AF) estimates derived from Illumina and Oxford Nanopore Technologies (ONT) sequencing analyzed with CleanFinder. (A) Correlation of genotype frequencies between Illumina and ONT (super accuracy basecalling mode) in bulk-edited samples (left) and single-cell–derived clonal lines (right). Pearson correlation coefficients are indicated. Insets highlight lower-frequency variants. Overall concordance between platforms is high, although individual outliers are observed, and with lower concordance for lower abundant genotypes (up to 25%, inset). (B) Representative example of homopolymer-associated indel miscalling in ONT reads from a Cas-edited HIF1A clone in human induced pluripotent stem cells (iPSCs). The gRNA target site is indicated, and a homopolymeric poly-A nucleotide stretch is highlighted. Small deletions detected within this region reflect ONT-specific sequencing and basecalling errors rather than true editing events.

**Figure S3.**
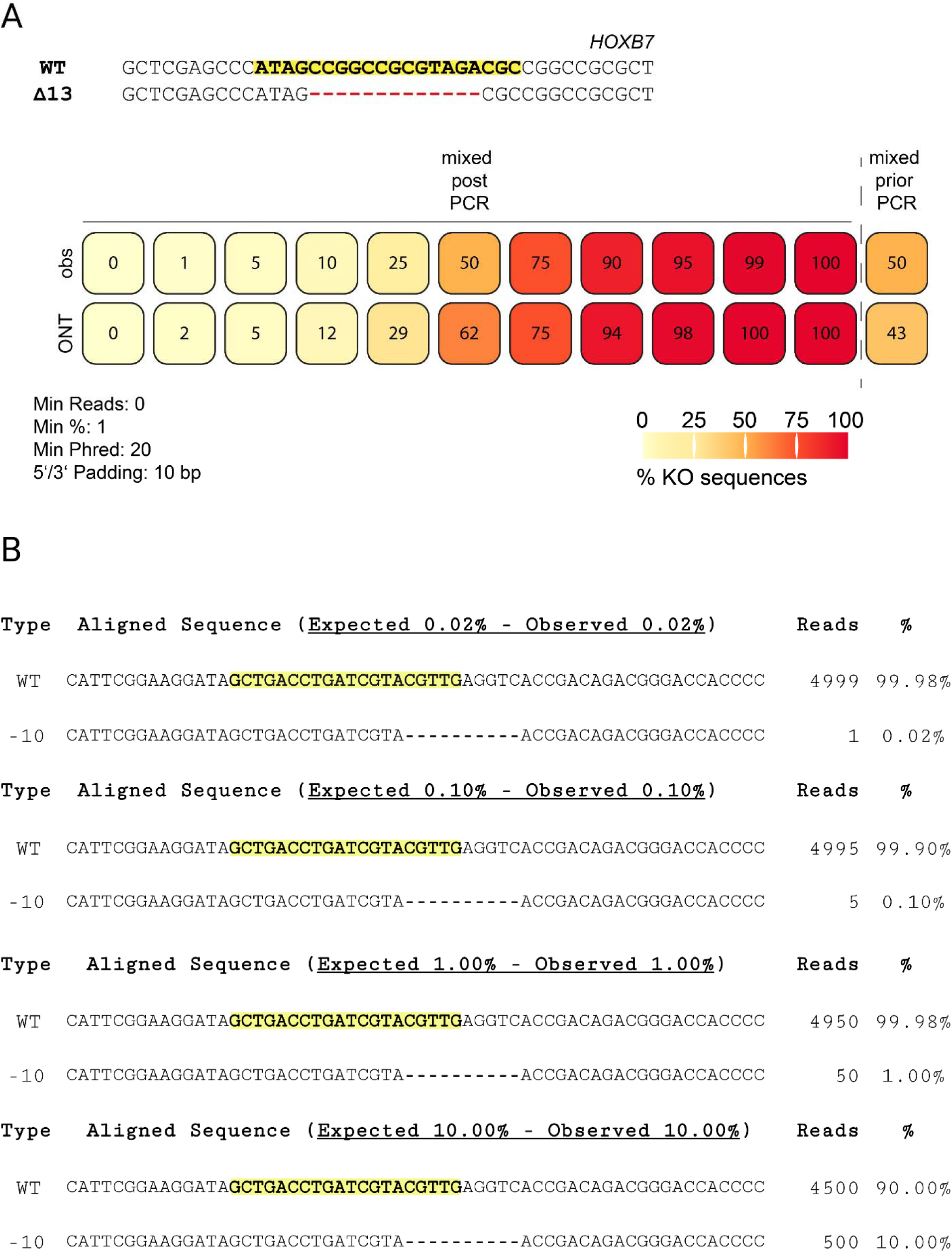
CleanFinder Sensitivity Assessment. (A) Experimental ONT sequencing of the *HOXB7* locus. Wild-type (WT) and Δ13 amplicons were mixed at defined proportions (0–100% KO), and allele frequency was quantified from Oxford Nanopore Technologies (ONT) reads. Colored boxes display the percentage of Δ13 (KO) reads detected at each input proportion. A 50% mixture generated prior to PCR is shown at the far right. Analysis parameters are indicated at the bottom (minimum reads = 0; minimum allele frequency threshold = 1%; minimum Phred score = 20; 5′/3′ padding = 10 bp). (B) In silico–generated Illumina-like reads. Defined proportions of WT and Δ10 sequences (0–100%) are shown across the top. Expected KO frequencies of 0.02%, 0.1%, 1%, and 10% were simulated and compared to the observed percentages after analysis.

**Figure S4.**
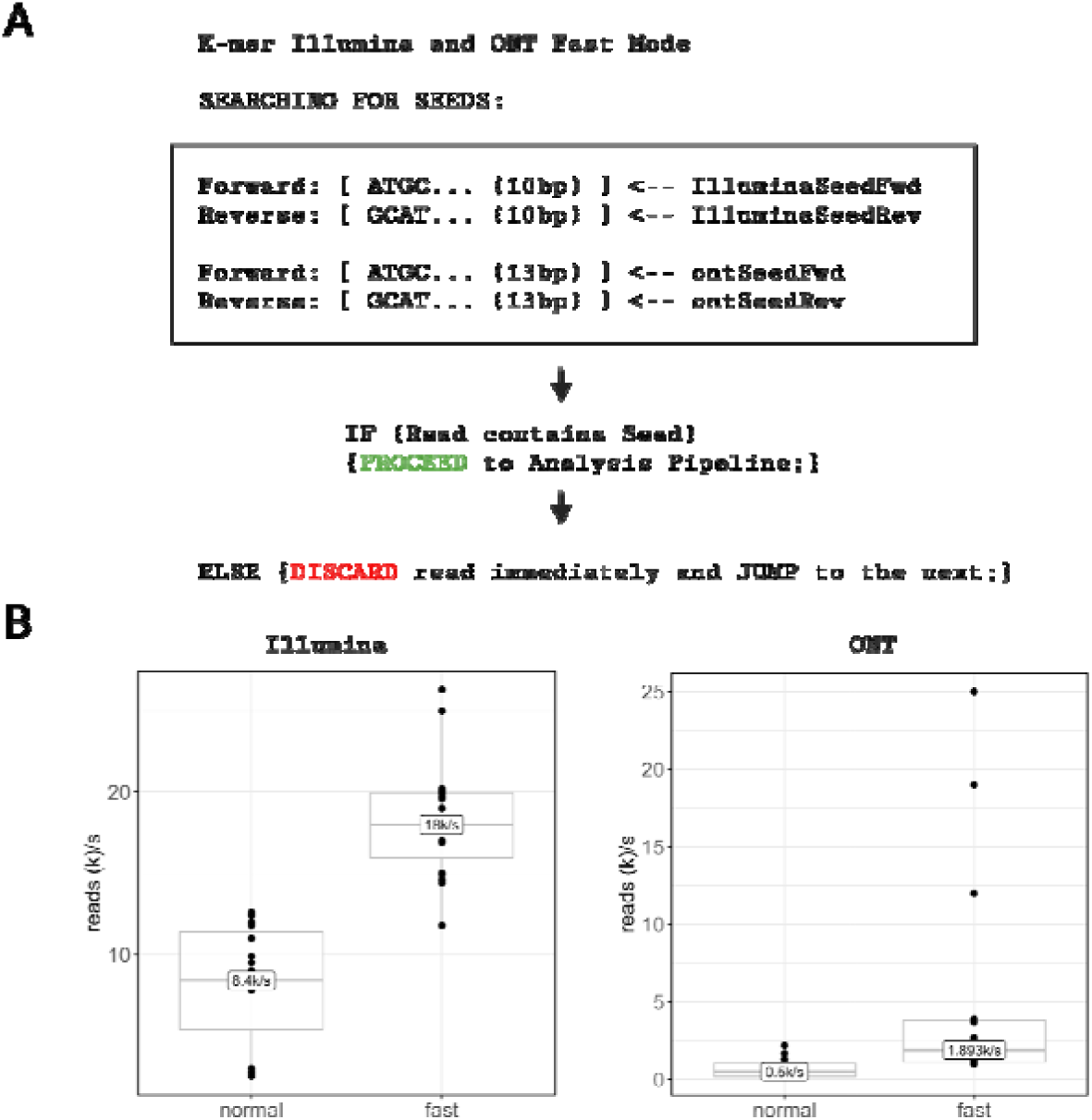
k-mer–guided pre-alignment filtering accelerates genome editing analysis. (A) Schematic representation of k-mer–based read filtering. Reads containing a target-specific k-mer seed proceed to full semi-global alignment, whereas reads lacking the seed are discarded prior to dynami programming. (B, left) Processing speed (reads per second, k/s) for Illumina datasets in normal alignment mode versus k-mer–accelerated fast mode. Median throughput increased from approximately 8.4k reads/s to 18k reads/s. (B, right): Processing speed for ONT datasets in normal versus fast mode. Median throughput increased from approximately 0.5k reads/s to 1.89k reads/s. Speed estimates were obtained from representative datasets and depend on file characteristics, including read length, dataset size, and the proportion of locus-unrelated reads. Datasets containing large fractions of unrelated or multiplexed reads benefit most from early k-mer–based filtering.

**Figure S5:**
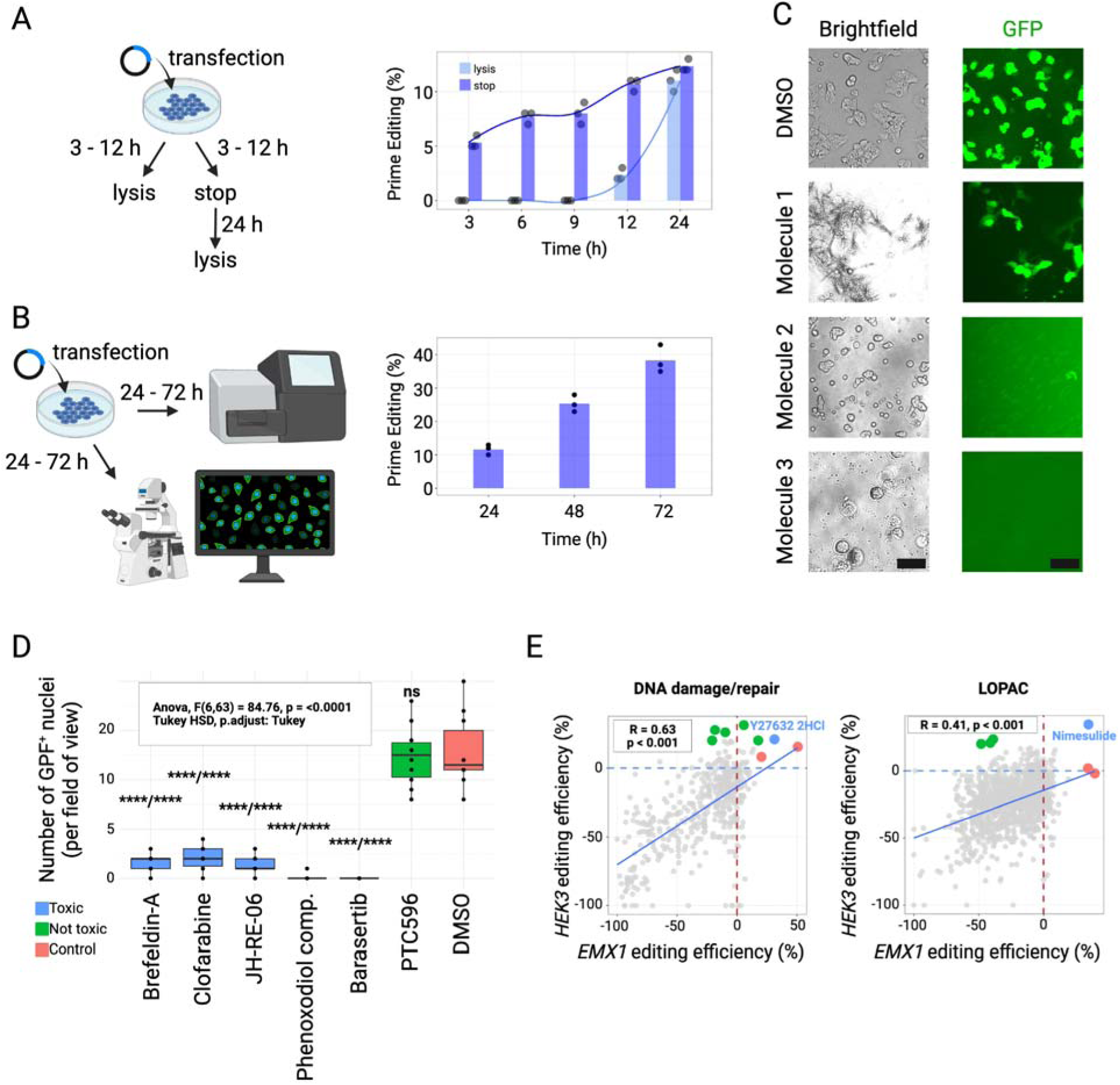
Time-resolved validation and screening-scale identification of prime editing modulators. (A) Schematic representation of the early time-course experiment. Cells were transfected and either lysed at early time points (3–12 h) or transfection was stopped at indicated time points and lysed 24h later.Quantification of prime editing efficiency over time shows progressive accumulation of edited alleles, with minimal signal at early time points and increasing editing detectable at later intervals. Bars represent mean editing efficiency, and dots indicate individual replicates. (B) Extended time-course analysis following transfection. Cells were analyzed at 24, 48, and 72 h post-transfection by fluorescence-based readout. Prime editing efficiency increased over time, demonstrating continued accumulation of edited alleles during the post-transfection period. (C) Representative brightfield and GFP fluorescence images under control (DMSO) and compound-treated conditions. DMSO-treated cells display robust GFP signals consistent with efficient editing. Molecule 1 shows reduced GFP-positive cells, indicating inhibitory activity, whereas Molecules 2 (no signal) and 3 (cytotoxic) exhibit differential modulation of GFP expression. Images were acquired 48 hr post-transfection. Scale bars = 200 µm. (**D**) Representative compounds exerting cytotoxic effects during the primary molecule screen. GFP-positive nuclei were counted in n = 10 independent fields of view and differences between compounds were assessed using Analysis of Variance (ANOVA) analysis with Tukey HSD post-hoc correction. Statistical significance was inferred at p < 0.05 and is annotated by asterisks versus: non-toxic compound/DMSO. ****: p < 0.001. (**E**) Scatter plots showing fold changes of prime editing efficiency at EMX1 (x-axis) and HEK3 (y-axis) loci following treatment with compounds from the Selleck (DNA damage/repair) and Sigma (LOPAC) libraries. Each dot represents a compound. Colored points indicate enhancers specific to EMX1 (green), HEK3 (red), or common to both loci (blue). Dotted lines represent DMSO baseline controls. Correlation analysis of prime editing efficiencies between *EMX1* and *HEK3* loci were calculated to indicate common trends regardless of base substitution and insertion. For this analysis, compounds with transfection issues were disregarded. Pearson’s correlation coefficient R is reported.

**Figure S6.**
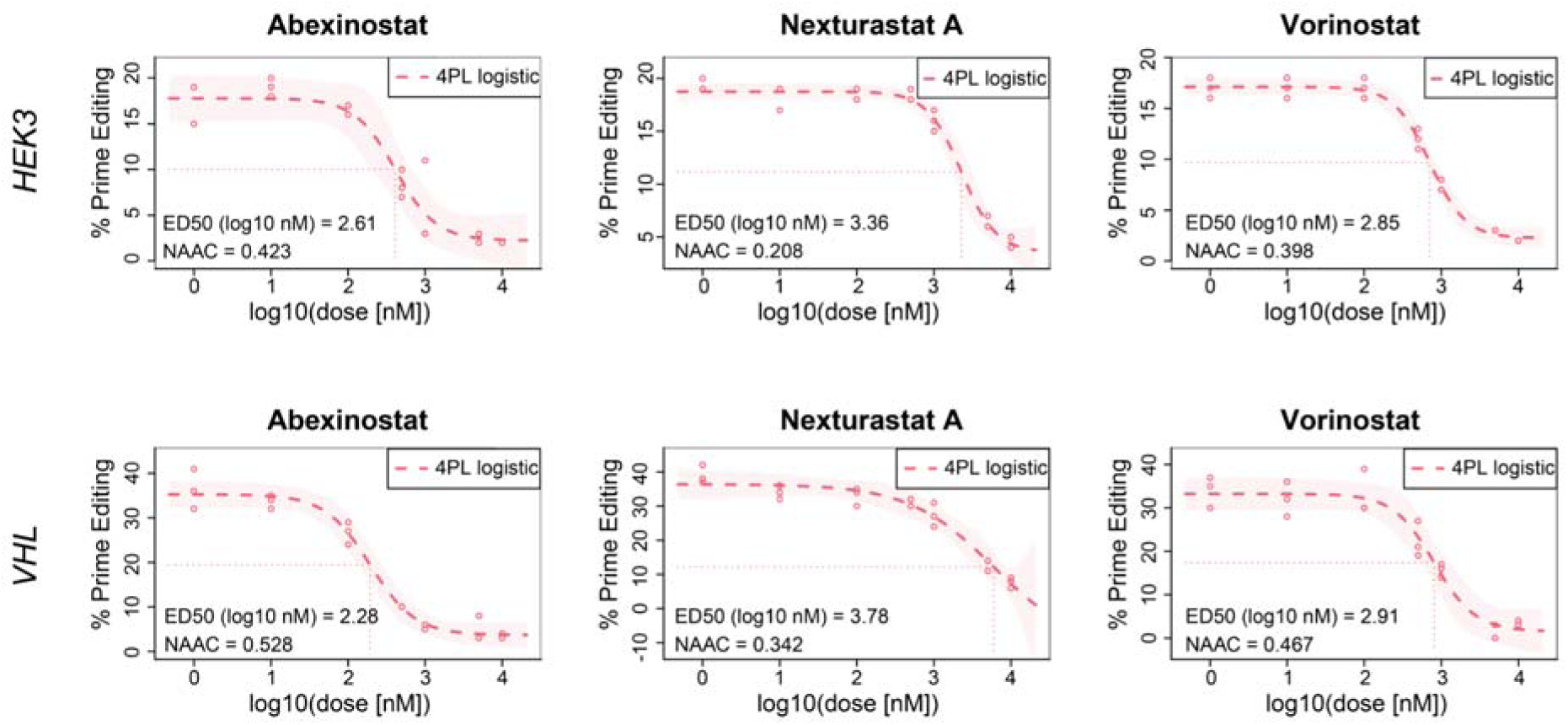
Supplementary Figure 6: Dose–response validation of HDAC inhibitor effects on prime editing across independent loci. Effect of HDAC inhibition on prime editing efficiency on base insertions for two distinct genetic loci, *HEK3* and *VHL*. Dose-response curves were calculated based on seven different concentrations of inhibitors in DMSO. 4 Parameter Logistic (4PL) was applied to model effects of HDAC inhibitors. ED50 representing the 50% effective dose, and normalized area above the curve (NAAC) values are indicated in the dose response plots.

**Figure S7.**
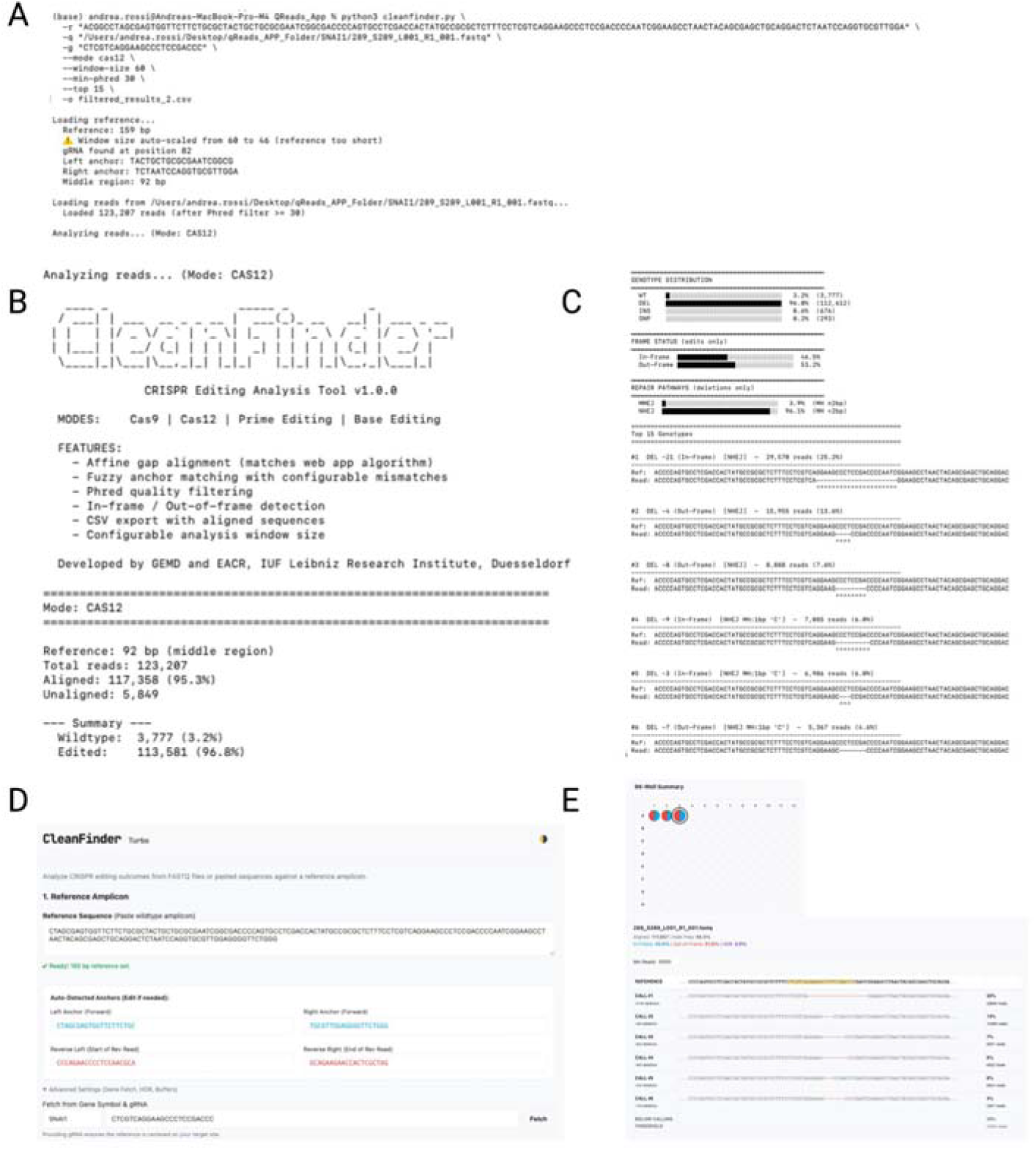
Implementation of CleanFinder as a Python CLI and Turbo browser module. (A) Example execution of the CleanFinder Python command-line interface (CLI). The reference amplicon is loaded, anchors are determined automatically, and analysis is initiated in the selected editing mode. Summary statistics report total reads, aligned reads, and editing fraction. (B) CLI header and feature overview indicating supported editing modalities (Cas9, Cas12, Prime Editing, Base Editing) and configurable analysis parameters including anchor length, mismatch tolerance, and optional gRNA-centered windowing. (C) Representative CLI output showing genotype distribution, frame classification (in-frame versus out-of-frame), and aligned top allele calls. The CLI additionally reports deletion-associated microhomology metrics for repair pathway annotation. (D) CleanFinder Turbo browser interface. The reference amplicon is pasted directly into the interface, and forward anchors together with reverse-complement anchors are automatically extracted. Users may configure mismatch tolerance and optionally provide HDR templates or gRNA sequences. (E) 96-well plate visualization in Turbo mode. Each well is displayed as a compositional map reflecting allele categories (wildtype, HDR, in-frame indel, out-of-frame indel, substitution). Selection of a well reveals allele-level alignment centered on the editing locus. Turbo performs heuristic anchor-guided classification for rapid screening and reports a reduced set of alignment-derived annotations compared to the full CLI implementation.

**Figure S8.**
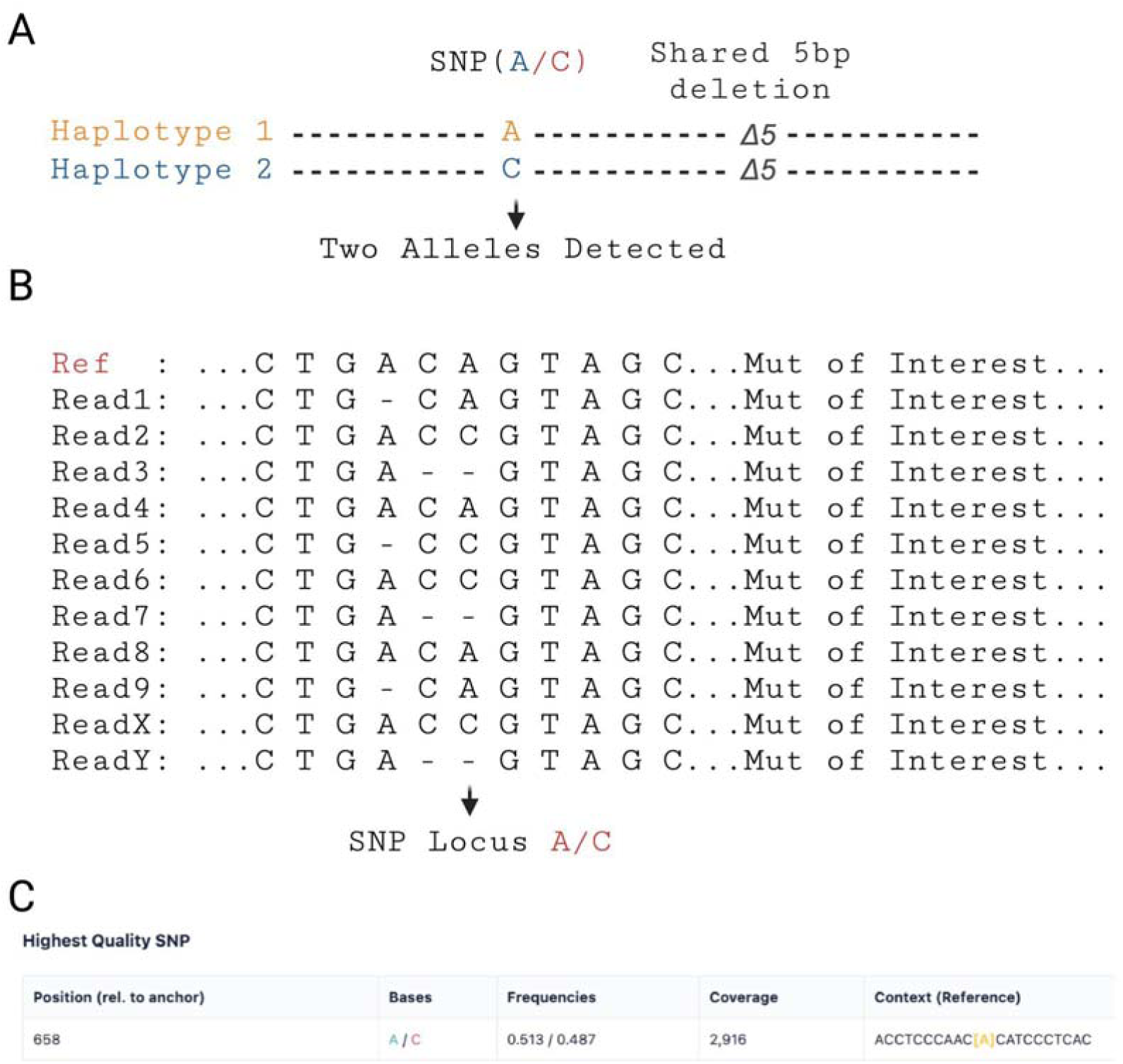
Detection of allelic dropout using heterozygous SNP tracking. **(A)** Conceptual representation of two parental haplotypes carrying a heterozygous SNP (A/C) and a shared 5-bp deletion introduced by CRISPR editing. Both haplotypes are detected within the sequencing reads, confirming the presence of two distinct alleles in the sample. (B) Schematic of the global alignment of long-read sequences (ONT) to the reference. Reads are aligned using a Needleman–Wunsch algorithm tolerant to insertions and deletions. Gaps (–) correspond to indels typical of ONT data but do not affect SNP counting. The SNP locus (A/C) is highlighted at the aligned position where base frequencies are calculated across all reads. (C) Example output of the *XPA* locus from the Allelic Dropout module. The table summarizes the highest-confidence heterozygous SNP identified within the amplicon, including its position relative to the anchor, detected bases, allele frequencies, read coverage, and reference sequence context. The near-equal frequencies (A = 0.513, C = 0.487) confirm a biallelic configuration, indicating no allelic dropout.

## Notes

https://github.com/andrearossi-lab/CleanFinder/tree/main

https://data.mendeley.com/datasets/54m5svs2rc/1

### Competing Interest Statement

The authors have declared no competing interest.

### Summary of Updates

The whole manuscript has been updated, with new alghoritm, data, and modules.

